# Altered structure and stability of bat-prey interaction networks in logged tropical forests revealed by metabarcoding

**DOI:** 10.1101/2020.03.20.000331

**Authors:** David R. Hemprich-Bennett, Victoria A. Kemp, Joshua Blackman, Matthew J. Struebig, Owen T. Lewis, Stephen J. Rossiter, Elizabeth L. Clare

## Abstract

Habitat degradation is pervasive across the tropics and is particularly acute in Southeast Asia, with major implications for biodiversity. Much research has addressed the impact of degradation on species diversity; however, little is known about how ecological interactions are altered, including those that constitute important ecosystem functions such as pest consumption.
We examined how rainforest degradation alters trophic interaction networks linking insectivorous bats and their prey. We used DNA metabarcoding to study the diets of forest-dwelling insectivorous bat species, and compared bat-prey interaction networks between old growth forest and forest degraded by logging in Sabah, Borneo.
We predicted that rainforest degradation would cause measurable reductions in the numbers of prey consumed by individual bats, and that this degradation would yield networks in logged forest with lower functional complementarity, modularity and nestedness than those in old growth forest.
Compared to bats in old growth rainforest, bats in logged sites consumed a lower diversity of prey. Their interaction networks were less nested and had a more modular structure in which bat species had lower closeness centrality scores than in old growth forest. These network structures were associated with reduced network redundancy and thus increased vulnerability to perturbations in logged forests.
Our results show how ecological interactions change between old growth and logged forests, with potentially negative implications for ecosystem function and network stability. We also highlight the potential importance of insectivorous bats in consuming invertebrate pests.

**Malay abstract:**

1. Degradasi habitat merupakan suatu fenomena yang berleluasa dikawasan tropika, terutamanya di Asia Tenggara dengan implikasi yang besar ke atas biodiversiti. Banyak kajian telahpun meneliti impak degradasi habitat atas kepelbagaian spesis. Walau bagaimanapun, dari segi mana interaksi ekologi diubah suai kurang diselidik, termasuk interaksi yang membentuk fungsi ekosistem yang penting seperti pemakanan binatang perosak.
2. Kami telah memeriksa bagaimana degradasi hutan hujan tropika dapat mengubah suai interaksi antara tahap trofik yang menghubungkan kelawar yang memakan serangga dan mangsa mereka. Kami telah menggunakan “DNA metabarcoding” untuk mengenal pasti kandungan artropod dalam sampel najis kelawar and membandingkan jaringan interaksi kelawar dan mangsa mereka diantara hutan dara dan hutan yang telah dibalak di Sabah, Borneo.
3. Kami meramalkan bahawa degradasi hutan hujan akan menyebabkan kekurangan dalam bilangan nod mangsa yang dimakan oleh setiap individu kelawar yang dapat diukur. Degradasi ini pula boleh menghasilkan jaringan yang mempunyai fungsi saling melengkapi dan modulariti yang rendah, dan lebih berkelompok atau “mempunyai “nestedness” yang lebih tinggi di hutan yang dibalak berbanding hutan dara.
4. Kelawar di kawasan hutan yang dibalak memakan diversiti mangsa yang lebih rendah dengan kelawar di habitat hutan hujan dara. Jaringan-jaringan interaksi mereka kurang berkelompok dan mempunyai stuktur yang lebih modular dimana spesis kelawar mempunyai pemarkahan kerapatan berpusat yang lebih rendah daripada sepesis kelawar di hutan dara. Struktur-struktur jaringan ini berkait dengan lebihan jaringan atau “network redundancy” yang lebih rendah and ini membawa kepada kerentantan yang meningkat terhadap gangguan luar di hutan yang telah dibalak.
5. Keputusan kami menunjukkan bagaimana interaksi ekologi berubah diantara hutan dara dan hutan yang dibalak, dengan potensi implikasi negatif untuk fungsi ekosistem dan kestabilan jaringan. Kami juga telah menunjukkan potensi kepentingan kelawar yang memakan serangga dalam fungsi mereka untuk makan perosak invertebrat.

**Data Accessibility Statement:** Data are currently archived at the Centre for Ecology and Hydrology Environmental Information Data Centre (https://doi.org/10.5285/8b106445-d8e0-482c-b517-5a372a09dc91) and will be released from embargo following publication. Specific analysis scripts are available on GitHub with links given in the manuscript and will be archived on Zenodo prior to publication.

**Statement of authorship:** SR, EC, DHB, MS and OTL conceived the project, DHB, VK and JB undertook field collections and laboratory work, DHB analysed the data with input from EC, and DHB wrote the manuscript with input from all authors.

## Introduction

Many tropical forests have been degraded by human activity, leading to biodiversity loss (Barlow et al., 2016) and modifying the ecological processes fundamental to forest dynamics (Ghazoul, Burivalova, Garcia-Ulloa, & King, 2015) such as the regeneration of plant communities. Land-use change is responsible for 62% of habitat alteration in Asia (Song et al., 2018), with degraded forests being of particular conservation interest; these habitats may retain high biodiversity yet have minimal protection and are vulnerable to clearance for agriculture and development (Meijaard et al., 2018).

The island of Borneo hosts high biodiversity but has lost much of its old growth forest, with 46% of its remaining forest classified as degraded by selective logging (Gaveau et al., 2014). As a consequence, there is considerable interest in understanding the conservation value and viability of these remaining forest areas (Meijaard & Sheil, 2007; Melo, Arroyo-Rodríguez, Fahrig, Martínez-Ramos, & Tabarelli, 2013), especially given their minimal conservation protection under current policies (Struebig et al., 2015). Mounting evidence suggests selectively-logged forests can support a substantial proportion of the original biota, and are generally more biodiverse than secondary forests (Gibson et al., 2011). Indeed, despite their degradation, Borneo’s logged forests retain potentially important communities of groundd-welling mammals (Deere et al., 2017), bats (Struebig et al., 2013), birds (Edwards et al., 2010) and invertebrates (Slade, Mann, & Lewis, 2011). Much less is understood, however, about how such habitat degradation impacts the ecological interactions among co-occurring species, such as between predators and prey, hosts and parasites, and plants and their pollinators (Andresen, Arroyo-Rodriguez, & Escobar, 2018).

A powerful approach for understanding ecological interactions is through network analyses, in which interactions (‘edges’) are represented by links among biological ‘nodes’ (usually species) (Cirtwill et al., 2018). These networks most commonly depict mutualisms such as pollination and seed dispersal (Bascompte, 2009) or antagonisms such as parasitism and predation (Lafferty, Dobson, & Kuris, 2006), quantifying aspects of the community’s trophic structure. Through measuring and comparing aspects of network structure, it is possible to predict a system’s resilience to perturbations (Memmott, Waser, & Price, 2004), the importance of a species to a given network function (Freeman, 1978), and the potential for competition between species and their conspecifics (Bastolla et al., 2009). Altered network structure may thus reveal functionally important shifts within ecological communities.

Highly mobile predators may be important for stabilising numbers of prey throughout their habitat (McCann, Rasmussen, & Umbanhowar, 2005; McCracken et al., 2012), by dampening boom and bust cycles of insects (Kunz, Torrez, Bauer, Lobova, & Fleming, 2011). Previously, lower bird abundance linked to forest degradation was shown to reduce top-down control of phytophagous herbivores, thus increasing herbivory (Peter, Berens, Grieve, & Farwig, 2015) and potentially affecting forest restoration (Böhm, Wells, & Kalko, 2011). Similarly, bats may control herbivorous insects in rainforests (Kalka, Smith, & Kalko, 2008). Therefore, the loss of bats may be expected to alter ecosystem functioning via trophic cascades.

Research in palaeotropical forests suggests logging affects bat community composition and abundance by altering roost availability (Struebig et al., 2013), reflecting patterns seen in the neotropics (Peters, Malcolm, & Zimmerman, 2006). While these communities might be predicted to show altered network structures, studies from mutualistic neotropical systems of bats dispersing seeds have shown little difference in network structure in fragmented forest, despite a reduction in the number of food species consumed (Laurindo, Novaes, Vizentin-Bugoni, & Gregorin, 2019), possibly as a result of highly resilient bat species which are core to their networks. Bat-seed dispersal networks have been shown to be robust to secondary extinctions (Mello et al., 2011), but parallels between mutualistic and antagonistic networks may be limited due to known differences in their structure (Lewinsohn, Prado, Jordano, Bascompte, & Olesen, 2006; Thébault & Fontaine, 2010). Therefore, given the key predation roles of insectivorous bats in rainforests, an improved understanding of their feeding ecology is a priority for the conservation of bats and their ecosystems (Meyer, Struebig, & Willig, 2016).

Genetic tools, particularly high throughput sequencing (HTS), are increasingly used for dietary analyses (Aizpurua et al., 2018; Clare, Fraser, Braid, Fenton, & Hebert, 2009; Czenze et al., 2018; Razgour et al., 2011). The application of DNA metabarcoding to bat and bird faeces makes it possible to obtain detailed information on previously unknown species interactions (Clare, 2014; Creer et al., 2016; Evans, Kitson, Lunt, Straw, & Pocock, 2016; Roslin & Majaneva, 2016). While traditional approaches based solely on the morphological identification of prey items in guano restricted the resolution of diet, metabarcoding approaches can allow numerous prey species to be identified at genus- or family-level (Clare, 2014), so providing the means to compare datasets of ecological interactions across networks.

Here we use DNA metabarcoding to assess the impact of rainforest degradation on predator-prey interactions, focusing on insectivorous bats that forage under the forest canopy in Borneo. We captured bats in old growth and logged rainforest and generated bipartite ecological networks of their interactions with prey using metabarcoding of their guano. Comparing the taxonomic composition, completeness and structure of these networks, we predicted that:

1. Disturbance causes the network in logged forest to have lower functional complementarity, modularity and nestedness than networks in old growth forest.
2. Bats in logged forest consume fewer prey items than in old growth forest, leading to higher closeness centrality in logged forests.

In addition, we screened the resulting sequence data for economically important pests of forestry plantations and agricultural crops in modified tropical landscapes.

## Methods

### Sample collection

We sampled bats using six harp traps per night at three sites in lowland tropical rainforest in Sabah, Malaysian Borneo, each <500m above sea level and experiencing a largely unseasonal climate. In total we sampled at 636 unique trapping locations over 876 trap nights. We collected faecal samples in two old growth sites: Danum Valley Conservation Area (hereafter ‘Danum’), Maliau Basin Conservation Area (‘Maliau’), and a forest heavily disturbed by multiple rounds of logging: the Stability of Altered Forest Ecosystems Project (‘SAFE’). Bats were captured by placing harp traps at regular intervals (mean 37m SD 77m) along landscape features such as streams and trails. The traps were erected in the morning, and then checked at approximately 8PM and 8AM. Bats were released at the points of capture, with pregnant, lactating or juvenile bats being released instantly. Otherwise, captured bats were placed into individual cloth bags for up to 12 hours, upon which any guano was removed and stored at −20°C. For full information on fieldwork see Supplementary Information 1.

### Laboratory work

To build a network of bat-insect interactions for each of the three forest sites studied (Danum, Maliau and SAFE), we sequenced prey DNA from bat guano using metabarcoding. DNA extraction, PCR, sequencing and quality-control took place following the methods outlined by Czenze et al. (2018). Briefly, we extracted DNA using a Qiagen stool kit, then amplified it using arthropod-specific primers (Zeale, Butlin, Barker, Lees, & Jones, 2011) and sequenced the DNA on an Illumina MiSeq (Supplementary Information 2).

### Bioinformatics

The resulting sequences were merged into contiguous reads, the primers were removed, and the reads were length-filtered and collapsed to haplotype with any singletons excluded from the resulting dataset, before clustering sequences into Molecular Operational Taxonomic Units (MOTUs) using the Uclust algorithm (Edgar, 2010) in QIIME (Caporaso, Kuczynski, Stombaugh, Bittinger, & Bushman, 2010). To reduce costs, we restricted sequencing to the ten bat species for which we were able to obtain at least ten guano samples from one or more forest sites (see Table 1 for sample sizes). This approach was taken to ensure that, as much as is possible, networks contained the same sets of bat taxa. Removing rare or unevenly distributed species was suggested by Blüthgen (2010) to reduce the confounding impact of observation frequency. We only took this approach for bats and not for MOTUs due to the expected comparative rarity of most MOTUs consumed.

**Table 1.** Bat species and samples used to construct the ecological networks.

|  | *Old growth forest* |  | *Logged forest* |
| --- | --- | --- | --- |
|  | *Danum* | *Maliau* | *SAFE* |
| *Hipposideros cervinus* (Fawn Roundleaf bat) | 184 | 90 | 110 |
| *Hipposideros diadema* (Diadem roundleaf bat) | 2 | 10 | 3 |
| *Hipposideros dyacorum* (Dayak roundleaf bat) | 0 | 13 | 9 |
| *Hipposideros ridleyi* (Ridley's roundleaf bat) | 2 | 1 | 14 |
| *Kerivoula hardwickii* (Hardwicke's woolly bat) | 3 | 0 | 23 |
| *Kerivoula intermedia* (Small woolly bat) | 29 | 9 | 44 |
| *Kerivoula papillosa* (Papillose wooly bat) | 21 | 0 | 6 |
| *Rhinolophus borneensis* (Bornean horseshoe bat) | 1 | 26 | 10 |
| *Rhinolophus sedulus* (Lesser woolly horseshoe bat) | 10 | 4 | 14 |
| *Rhinolophus trifoliatus* (Trefoil horseshoe bat) | 14 | 19 | 28 |

Where not otherwise stated, we generated the three networks by clustering sequences into MOTUs at 0.95 similarity, chosen to balance over- and under-splitting of MOTUs. We then compared representative sequences of each MOTU to one another using BLAST+ (Camacho et al., 2009), with the resulting data being filtered in LULU (Frøslev et al., 2017) to combine suspected duplicate MOTUs. However as the choice of clustering threshold used to cluster the sequence data into prey MOTUs can have a strong effect on the conclusions drawn (Clare et al., 2016; Hemprich-Bennett et al., 2018), we examine a range of clustering thresholds for a subset of the analyses in Prediction 1 to ensure that our conclusions are robust to our choice of this key parameter.

For a subset of analyses indicated below, networks were generated for each site at every MOTU clustering level from 0.91-0.98 similarity before quality control in LULU, allowing us to test the robustness of conclusions to changes in clustering level used.

### Analysis

We imported binary adjacency matrices generated into R version 3.4.4 (R Core Team, 2017) for analysis. For network-level analyses these matrices were then summed by bat species (i.e. *a_ij_* denotes all instances of bat species *i* consuming MOTU *j*), giving weighting to the network.

### Prediction 1: Disturbance causes the network in logged forest to have lower functional complementarity, modularity and nestedness than networks in old growth forest

To compare networks we focus on three measured components of network structure: nestedness, modularity and functional complementarity (Figure 1). Nestedness represents the extent to which the interactions of specialist nodes are nested subsets of the interactions of the generalist nodes (Almeida-Neto et al., 2008). Highly nested communities are more resilient to perturbations (Memmott et al., 2004), as the generalists and specialists perform the same role, conferring redundancy. Decreases in the nestedness of plant-pollinator communities following disturbance leads to reduced functional redundancy (Soares, Ferreira, & Lopes, 2017). We here calculate two metrics used to measure nestedness: discrepancy (Brualdi & Sanderson, 1999), and weighted nestedness based on overlap and decreasing fill (WNODF) (Almeida-Neto, Guimarães, Guimarães, Loyola, & Ulrich, 2008). Modularity is the extent to which a network’s interactions are partitioned into weakly-coupled ‘modules’ (Rezende, Albert, Fortuna, & Bascompte, 2009), which can contain the negative effects of perturbation (Fortuna et al., 2010). Modularity tends to decrease as prey availability is reduced (Oliveira, 2018), potentially increasing susceptibility to adverse effects of future stressors. Functional complementarity (Blüthgen & Klein, 2011; Devoto et al., 2012; Peralta, Frost, Rand, Didham, & Tylianakis, 2014), calculates the extent to which species have complementary non-overlapping diets by measuring the branch lengths of a functional dendrogram of their dietary dissimilarity. These metrics describe some of the most important elements of network structure and respond reliably to alterations to MOTU clustering level (Hemprich-Bennett et al., 2018), while allowing us to assess how phenomena such as habitat alteration affect ecosystem functioning.

**Figure 1).**
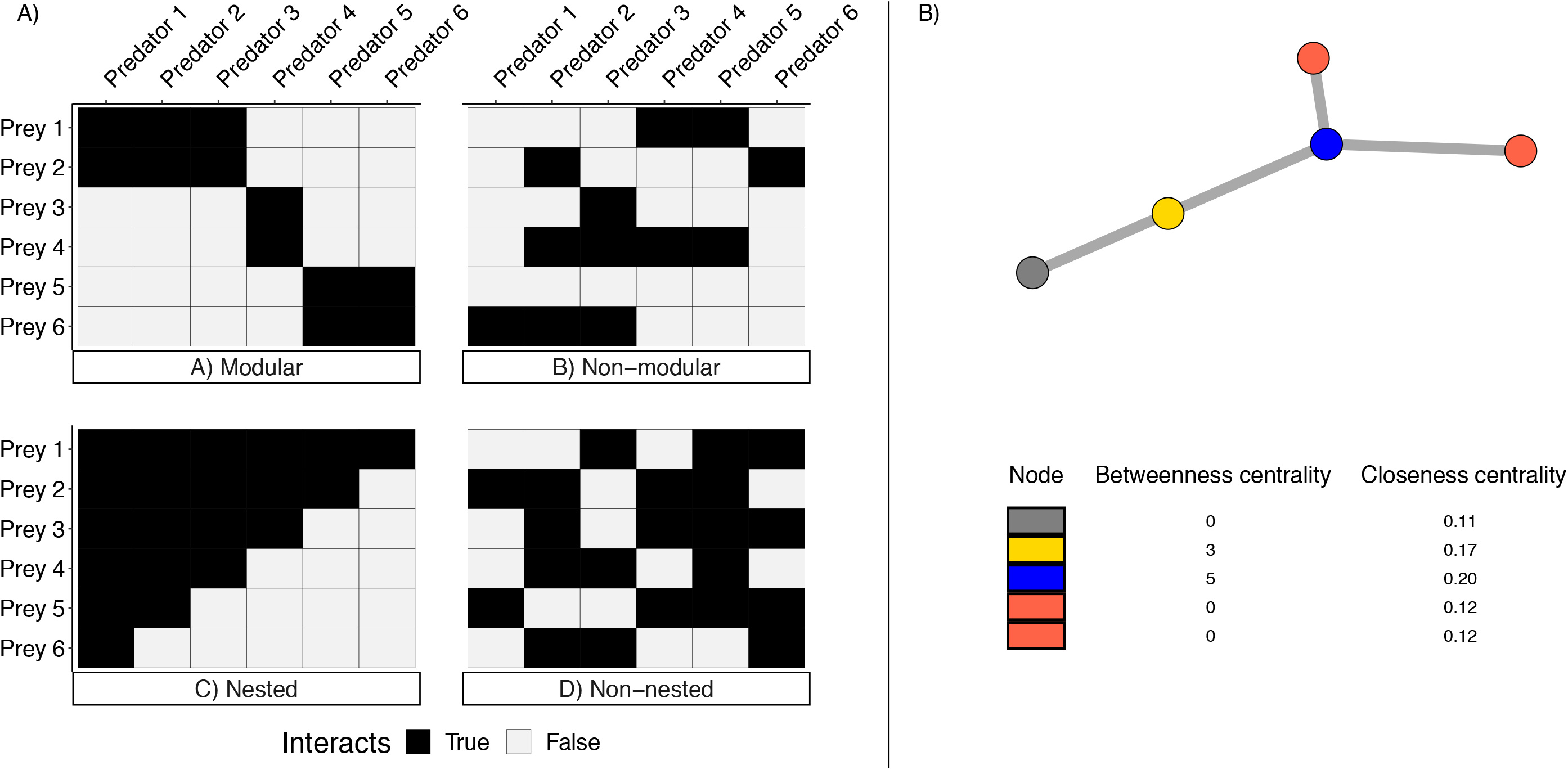
Panel 1 shows networks characterised by extreme a) modularity, b) non-modularity, c) nestedness structures and d) non-nestedness in bipartite networks. Panel 2 shows a simple network with values of betweenness centrality and closeness centrality for each node to 2 decimal places.

**Figure 2.**
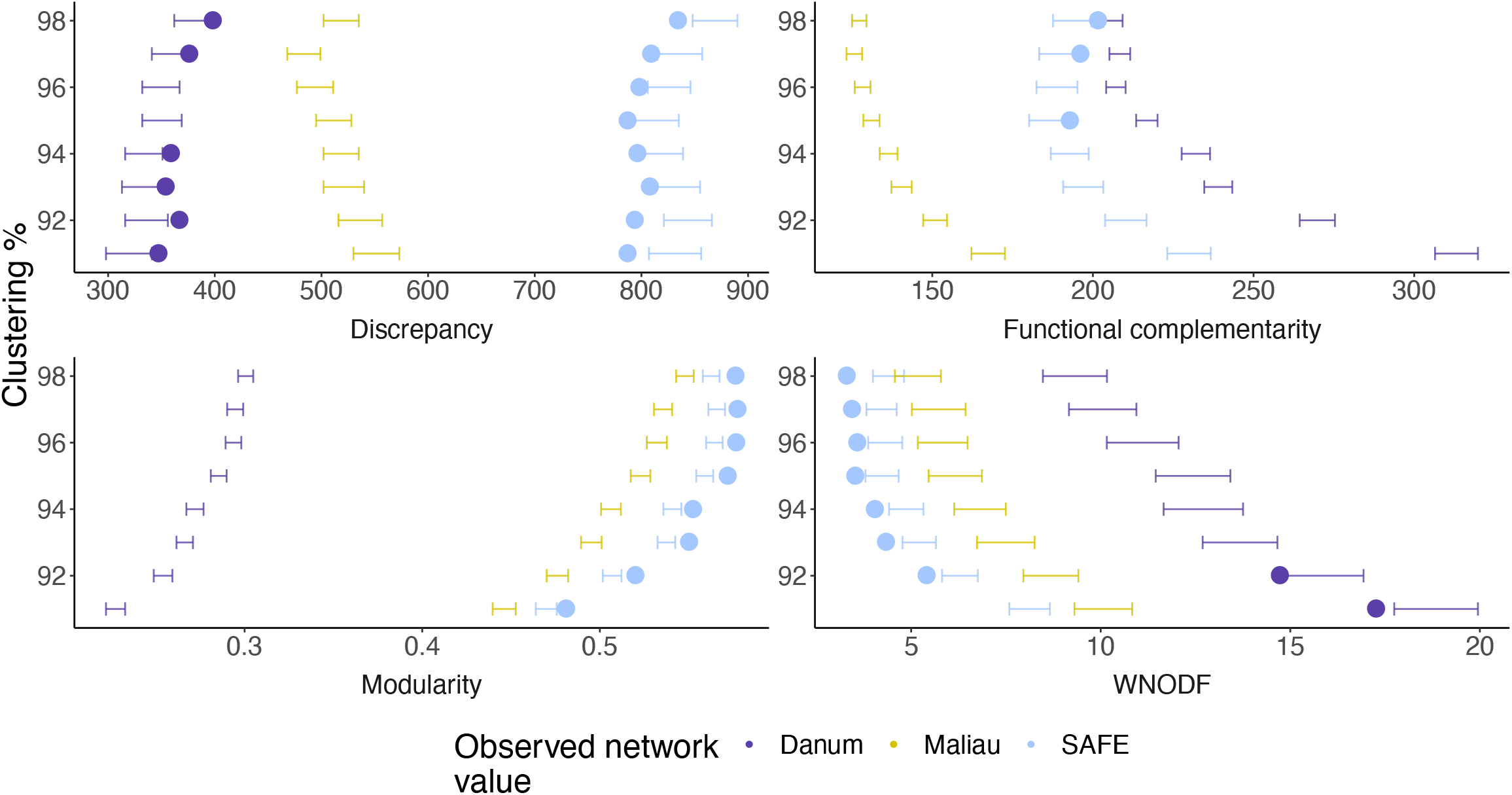
Summary plot showing the observed values (circles) and 95% confidence intervals (bars) given by the random values generated for each metric studied at each MOTU clustering level, showing how conclusions drawn are altered by MOTU clustering level. The observed values for each network were only plotted for the networks and metrics where the observed values fell outside of the range of 95% of the random values. For most metrics studied, the 95% confidence intervals do not overlap at most clustering levels used, showing that the networks differ regardless of clustering level used. Danum and Maliau are the old growth sites, and SAFE the logged site.

To test whether each of these metrics differ significantly between habitats more than would be expected by chance, we obtained null distributions for each metric, network and clustering threshold using the swap algorithm (Dormann & Strauss, 2014) to randomize each network for each MOTU clustering level 1,000 times, preserving the row and column sums. The observed value was deemed to be significantly different from chance if it was outside of the range of 2.5-97.5% of the randomly generated values. Two networks were also deemed different from one another if their ‘expected’ ranges did not overlap. Choice of MOTU clustering threshold in ecological metabarcoding studies has been shown to alter measurements of most network metrics (Hemprich-Bennett et al., 2018), and so to assess the impact of node resolution on the networks analysed here, we performed this analysis on data generated at each MOTU clustering threshold between 91-98% using the package ‘LOTUS’ (Hemprich-Bennett et al., 2018), which acts as a wrapper for the bipartite package (Dormann, Gruber, & Fründ, 2008). A conclusion can be considered to be robust if it is consistently found across all MOTU clustering thresholds used.

As sample size varied greatly across species and site (Table 1), we checked the impact of sample size and species diversity included in our analysed networks by using random subsamples of the bats captured at each site to generate smaller networks ranging from 40 individuals to the full network dimension, with 1,000 iterations per increment. Each focal metric other than modularity was calculated for the subnetworks, and the Shannon diversity (Shannon, 1948) of nodes used to create the network was recorded. These values were plotted to observe if network size (number of individuals used to make the network) or bat diversity were important determinants of network structure. If the rank order of a measured metric was not shown to be strongly determined by network size or bat diversity, then any conclusions drawn from it can be considered robust to sampling effort.

To determine the contribution of a given bat species to the measured networks, we also generated subnetworks by removing each species individually from the original networks and calculating each network metric. The influence of the species was then calculated by subtracting the subnetwork’s measured value from the whole-network value. We then ranked these calculated differences to show which species had the greatest and smallest impact on each network metric per site.

To obtain information on the taxonomic composition of bat diets, we compared a representative sequence for each MOTU using BLAST+ 2.7.1 (Camacho et al., 2009) against a database of arthropod CO1 sequences from the Barcode Of Life Database (BOLD) (Ratnasingham & Hebert, 2007), as accessed on 27/04/18. Using the program MEGAN 6.11.7 (Huson et al., 2016) and the quality-control parameters outlined in Salinas-Ramos et al., (2015), we excluded all sequences that could not be assigned to Order level, and used the BLAST assignments to determine the taxonomic composition of each guano sample. For each bat species at each site, we calculated the proportion of individuals that consumed a given Order of prey. We focussed on taxonomic Order (rather than, e.g. Family or Genus) due to the greater success in sequence assignment success at this level as sequence library completeness for Bornean arthropods is low.

### Prediction 2: Bats in logged forest consume fewer discrete prey items than in old growth forest, leading to higher closeness centrality in logged forests

We calculated the degree (number of prey MOTUs consumed) for each individual bat using the R package ‘bipartite’ (Dormann et al., 2008) and analysed these data with a fixed effects model, using species, habitat type (old growth or logged forest) and site as fixed effects, using backwards model selection with the Akaike information criterion (AIC), to detect whether models using habitat type or site were stronger predictors of bat degree. For each bat species we also calculated two measures of centrality using bipartite. Measures of centrality identify the influence of a node within a system or the distribution of its influences, often based on path lengths between nodes (Delmas et al., 2019). We focus on closeness centrality and betweenness centrality (Figure 1). Closeness centrality uses the shortest path lengths between all pairs of nodes to measure the proximity of the nodes in the network to one another, providing a measure of how rapidly a perturbation can spread (Freeman, 1978). If habitat degradation reduces the diversity or richness of prey available to predators, network metrics such as closeness centrality (Martín González, Dalsgaard, & Olesen, 2010) may increase as the nodes become ‘closer’ together. Betweenness centrality, in contrast, identifies the number of times a node is in the shortest path-length between any two other nodes, and so quantifies the importance of the node in connecting the overall network (Freeman, 1977). Using these measures of centrality, researchers have attempted to quantify the concept of ‘keystone species’ within the context of mutualistic networks (Martín González et al., 2010; Mello et al., 2015). In networks of frugivory, high centrality is linked both to the taxonomic class of a node, and the node having a high level of dietary specialisation (Mello et al., 2015), but in pollinators high centrality is associated with generalism (Martín González et al., 2010).

In addition, to assess the potential presence of prey species in bat diets, we compared our sequence data to publicly-available sequences on BOLD (Ratnasingham & Hebert, 2007) on 01/06/18 using the R package ‘bold’ (Chamberlain, 2019). We assigned sequences to species level using the highest obtained ‘similarity’ score per sequence if it was >0.98. The output data were then compared to a list of Malaysian crop pest species names obtained from Vun Khen (1996).

All code used for analyses in this paper can be found at: https://github.com/hemprichbennett/bat-diet; see Supplementary Information 2 for additional detail on laboratory work and bioinformatic analyses.

## Results

We captured 3,292 bats of 41 species, providing 700 faecal samples of 10 species that were used to create ecological networks (see Table 1). In total the 700 faecal samples yielded 18,737,930 contiguous reads, which were used to assemble the paired-end files. After removing adapters and primers, and any sequence with incomplete adapter or primer, this was reduced to 10,064,815 sequences, which was further reduced to 932,459 unique haplotypes after collapsing to haplotype, removing singletons, and discarding sequences outside of 2bp of the expected read-length. At 95% clustering this was condensed to 14,623 MOTUs, which LULU then reduced to 3,811 MOTUs (see Supplementary Information 3).

### Prediction 1: Disturbance causes networks in logged forest to have lower functional complementarity, modularity and nestedness than in old growth forest

Null models (Figure 3) indicated that the logged site was consistently less nested than the old growth sites (using the metrics discrepancy and WNODF). In an old growth forest site (Danum), values for functional complementarity were almost always within the expected range. Modularity was only significantly different from expectation in the logged forest, but it was always more modular than the old growth sites. No metrics analysed showed alterations in their rank order between the different MOTU clustering thresholds used, and so any conclusions drawn are unaltered by this bioinformatic parameter. Low values of discrepancy and high values of WNODF indicate a nested structure, a low value of modularity indicates a lack of modular structure, and a low value of functional complementarity indicates no complementarity between the predators.

**Figure 3.**
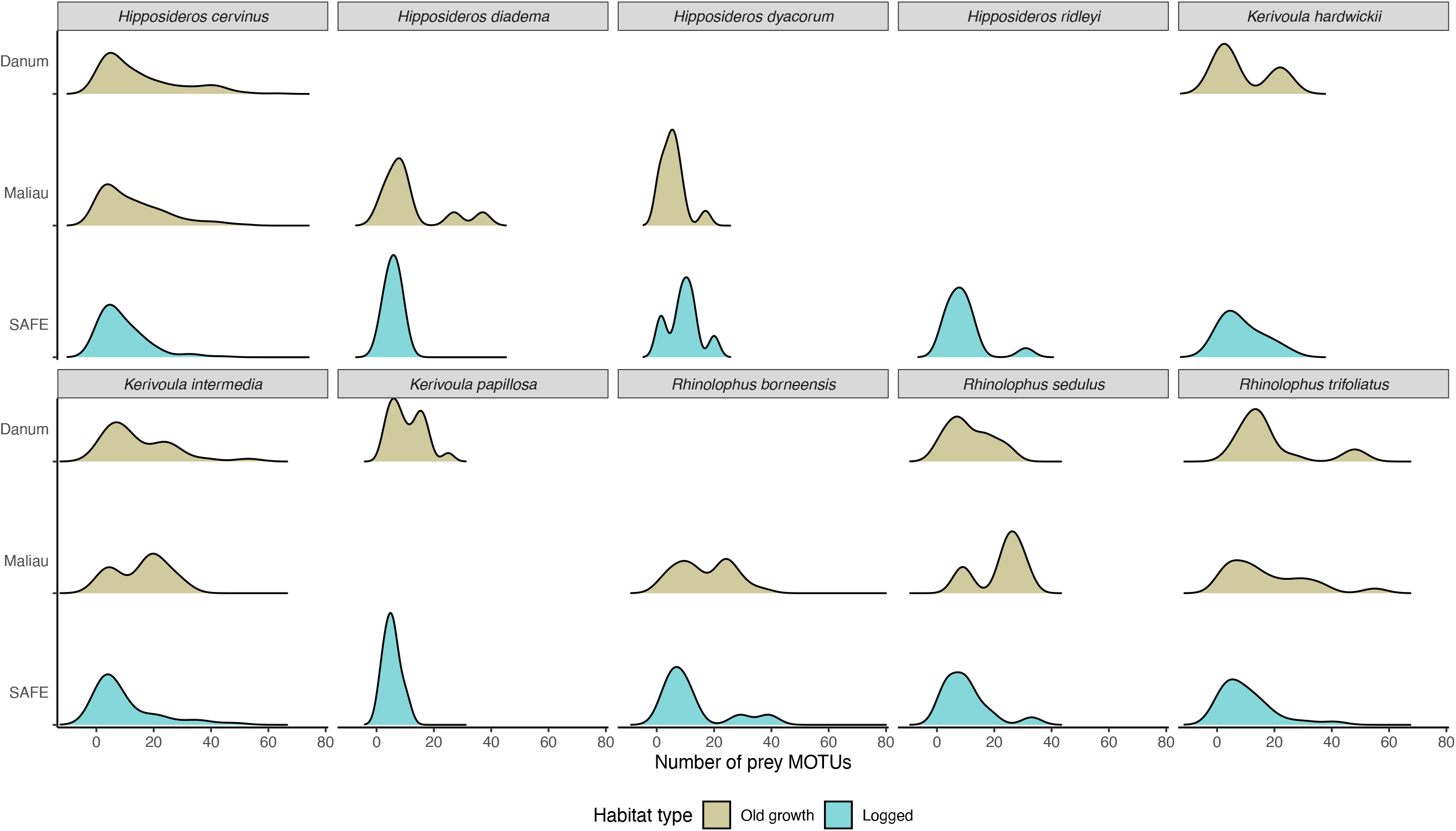
Smoothed histograms showing the number of MOTUs consumed by the individual bats for each focal bat species at each site. Species on average consumed a greater number of MOTUs in old growth forest than in logged forest.

Functional complementarity in logged forest was intermediate between the old growth sites, and likely not impacted by habitat degradation. Contrary to previous suggestions (Fründ, McCann, & Williams, 2016) we did not find that nestedness performed poorly with small sample sizes.

Most network metrics were greatly altered by bat removal, with the common species *H. cervinus* and *K. intermedia* causing the largest alteration to most metrics (see Supplementary Information 6). *R. borneensis* was shown to be important for the discrepancy, functional complementarity and modularity of an old growth site (Maliau).

Rarefaction revealed little impact of the diversity or richness of bats included in a network on any observed metric, but strong effects of the number of samples used to generate a subnetwork (see Supplementary Information 7). Discrepancy, functional complementarity and WNODF showed distinctions between logged and old growth forest sites once sampling effort approached completion.

### Prediction 2: Bats in logged forest consume fewer discrete prey items than in old growth forest, leading to higher closeness centrality in logged forests

We found a significant difference in degree for bats in old growth versus logged forest (F: 84.84 on 11 and 688 DF, p < 0.01, adjusted R^2^ = 0.57; see Table 2). The effect of habitat type on the number of MOTU consumed by an individual bat (its degree) was greater than the effect of species identity (Table 2, Figure 3), with bats in old growth forest consuming a greater number of MOTUs than bats in logged forest. The difference was lowest in *Hipposideros* species. This lower degree in logged forest shows that bats in this habitat generally consumed a lower number of prey items than their conspecifics in old growth rainforest.

**Table 2.** Degree model coefficients from fixed effects model, testing for the effects of habitat type (logged or old growth) and species identity on the degree of the individual bats studied.

| **Term** | **Estimate** | **Std error** | **Statistic** | **P value** |
| --- | --- | --- | --- | --- |
| Habitat: Logged | 9.078 | 0.874 | 10.384 | <0.001 |
| Habitat: Old growth | 14.330 | 0.630 | 22.740 | <0.001 |
| *Hipposideros diadema* | -2.680 | 2.948 | -0.909 | 0.364 |
| *Hipposideros dyacorum* | -5.045 | 2.458 | -2.053 | 0.040 |
| *Hipposideros ridleyi* | 2.407 | 2.820 | 0.854 | 0.394 |
| *Kerivoula hardwickii* | -0.915 | 2.336 | -0.392 | 0.695 |
| *Kerivoula intermedia* | 0.708 | 1.382 | 0.512 | 0.609 |
| *Kerivoula papillosa* | -3.681 | 2.230 | -1.651 | 0.099 |
| *Rhinolophus borneensis* | 4.198 | 1.928 | 2.178 | 0.030 |
| *Rhinolophus sedulus* | 0.617 | 2.201 | 0.281 | 0.779 |
| *Rhinolophus trifolius* | 2.491 | 1.552 | 1.605 | 0.109 |

All bat species had comparably low closeness centrality within the logged forest (see Supplementary Information 4), consistent with the observation that the logged forest had lower connectance than at the old growth forest sites. This shows that the bat species nodes were further from all other nodes in their network than in old growth rainforest. Rather than a reduced number of dietary items generating homogenous diets, this indicates the interactions of the network becoming more dispersed. At one of the old growth sites (Maliau) *Hipposideros cervinus* and *Rhinolophus borneensis* were the only species to have non-zero betweenness centrality scores (see Supplementary Information 5), indicating that every shortest path-length between nodes at Maliau (old growth) went via one of this pair of species, as opposed to the more diverse range of shortest path-lengths found in the other two networks.

Lepidoptera, Diptera (especially Cecidomyiidae) and Blattodea (especially Ectobiidae) were the most common prey Orders consumed (Figure 4; Supplementary Information 8). The lepidopteran pest species *Pleuroptya balteata* was detected in the diet of several bat species (Table 3) and at each site sampled, and *Psilogramma menephron* was consumed by *H. cervinus* in the logged forest site. However relatively few individual bats were recorded as consuming these species.

**Figure 4.**
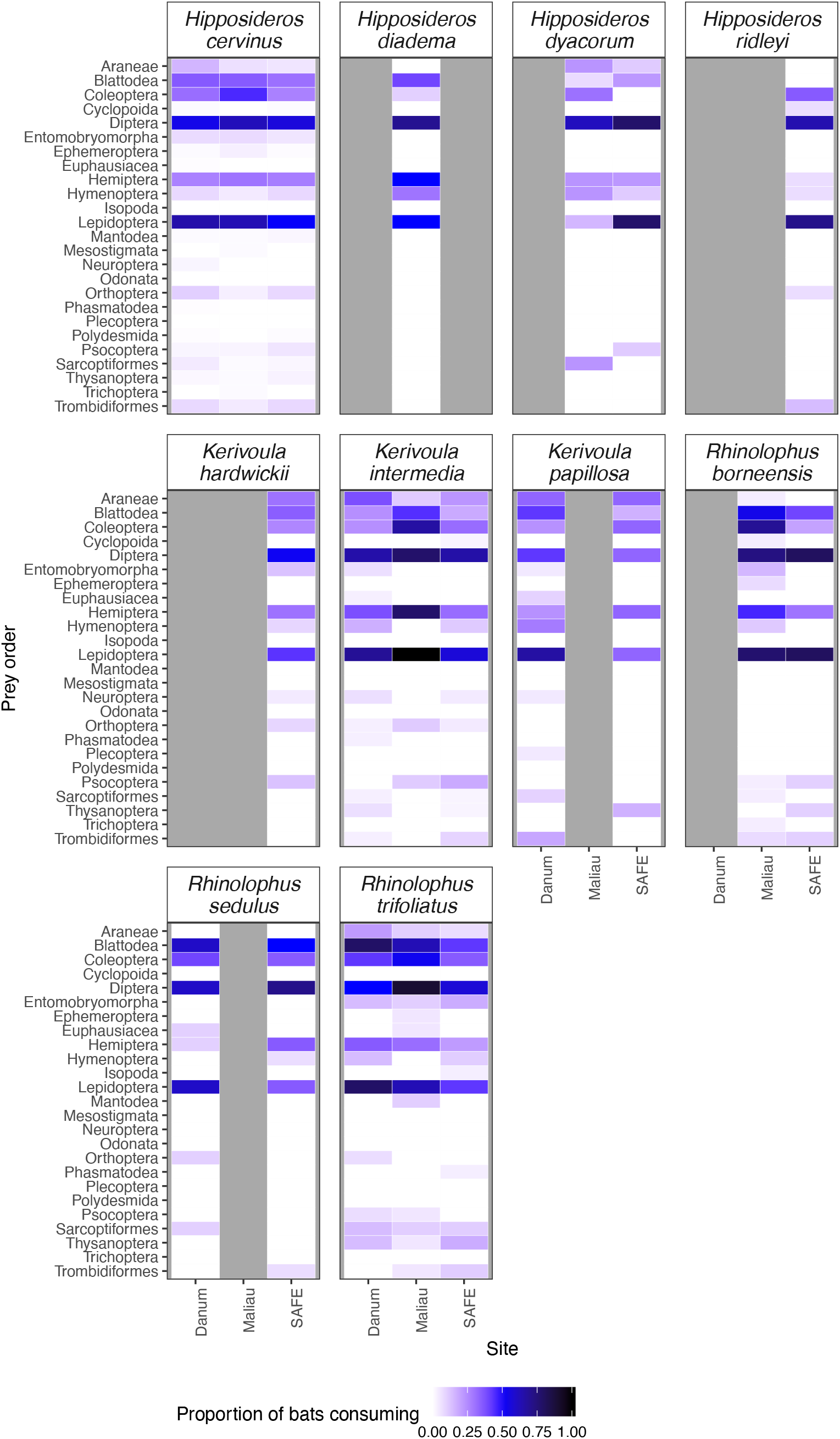
The proportion of individual bats of each species found to consume each taxonomic Order at each site studied.

**Table 3.** The crop pest species detected in the diet of bats at each of the study sites in Sabah.

| Pest species | Bat species | Site | Forest type | Number of detections /<br>number of bats sampled<br>at that site |
| --- | --- | --- | --- | --- |
| *P. balteata* | *H. cervinus* | Danum | Old growth | 3/184 |
| *P. balteata* | *H. diadema* | Danum | Old growth | 1/2 |
| *P. balteata* | *R. trifolius* | Danum | Old growth | 1/14 |
| *P. balteata* | *H. cervinus* | Maliau | Old growth | 2/90 |
| *P. balteata* | *H. cervinus* | Maliau | Old growth | 1/90 |
| *P. menephron* | *H. cervinus* | SAFE | Logged | 1/110 |
| *P. balteata* | *H. cervinus* | SAFE | Logged | 1/110 |
| *P. balteata* | *K. intermedia* | SAFE | Logged | 1/44 |
| *P. balteata* | *R. sedulus* | SAFE | Logged | 1/14 |
| *P. balteata* | *R. trifolius* | SAFE | Logged | 1/28 |

## Discussion

We found substantial differences in bat-insect interactions across sites experiencing varying degrees of habitat degradation. Bats consumed significantly fewer prey in logged forest sites than old growth forest; indeed, habitat type had a stronger effect on the number of MOTUs consumed by an individual bat than species identity.

Network structure also differed in several key aspects between the logged forest and old growth sites. Structural differences in centrality, modularity and nestedness together indicate that logged forest networks are more specialised than old growth rainforest networks. Systems that are specialised in this manner have been shown to be more vulnerable to extinctions than those with a more generalised structure (Memmott et al., 2004), such as the old growth rainforest networks analysed. Rainforests in Southeast Asia are facing multiple stressors, including the effects of disturbance, habitat fragmentation and climate change (Deere et al., 2020; Struebig et al., 2015). Our findings of altered network structure in an area which has been selectively logged indicate that such logged forests may be more sensitive to the effects of these future perturbations.

Bats foraging within the logged forest site consistently consumed fewer prey MOTUs than those within old growth forest. Indeed, the effect of habitat type was greater than that of the bat species in question, showing a strong alteration to foraging activity within logged forest. This was mirrored by the findings of reduced closeness centrality within the logged network, as the positions of bats within the network shifted. The most abundant bat species were found to have key roles in the structuring of their networks. Common predators will encounter a greater richness of prey than rarer species, through the ecological sampling effect (Dormann, Fründ, & Schaefer, 2017). While rare nodes are thought to have a stabilising effect on ecological networks (McCann, 2000) and are of conservation interest, abundant species are likely key to ecosystem functioning (Baker et al., 2018) as they are involved in a high proportion of the trophic energy transfer within a system. A possible strategy for conservation of ecological function could therefore be to find the species most important to a system and target them (Montoya, Rogers, & Memmott, 2012). If using this framework, we find that the most common bat (*Hipposideros cervinus*) is likely the species most key to the network, while also being the species with the least reduction in the number of prey MOTUs it consumes in the logged forest site (Figure 3).

Two species of moths known as crop or forestry pests were found in bats diets: *Pleuroptya balteata* was consumed at all sites and *Psilogramma menephron* was consumed in the logged forest. This is consistent with the potential role of tropical bats in pest control: although they represented only a small percentage (0.4%) of the MOTUs consumed by the bats overall, they were foraging in forest habitat and so the prey are likely occurring at lower densities than they would be expected to in managed landscapes. Natural habitats are thought to be important sources of pests to agricultural landscapes (Tscharntke et al., 2007) and so their consumption by predators is potentially of some economic importance; in this case *P. menephron* is an important pest of timber trees and *P. balteata* feeds on leaves of mango, tea and rambai (Vun Khen, 1996). In the neotropics (Kalka & Kalko, 2006; Kalka et al., 2008; Morrison & Lindell, 2012; Williams-Guillén, Perfecto, & Vandermeer, 2008) and temperate Europe (Böhm et al., 2011) bats are important agents controlling insect herbivory, but there are few examples from Southeast Asia (cf. Maas, Clough, & Tscharntke, 2013). We here provide one of few examples of bats in the region consuming pests, potentially reducing numbers of such species in natural habitats.

### Limitations

Due to the highly labour-intensive nature of capturing forest-interior bats it was only possible to sample three ecological networks. With limited replication (only one logged forest site and two old growth sites sampled), it is not possible to attribute differences between the sites unambiguously to the effects of logging, rather than other site-specific differences. Nonetheless, this work documents marked differences in network structure across the landscape that are consistent with variations in forest management, and which is likely to have implications for community stability and dynamics. It also highlights the utility of metabarcoding-based approaches for more comprehensive investigation of between-habitat differences in tropical forest predator-prey networks.

### Conclusions

Through combining DNA metabarcoding and network analysis, we have been able to measure how the ecological interactions which structure ecological communities differ between communities in logged and old growth forest. We show that in a logged forest bats and their prey exhibit altered network structures, which make them more prone to future local extinctions, adding to the previous findings that logged forest bat communities have altered composition and abundance (Struebig et al., 2013). Logged forests, although often heavily degraded, comprise a large proportion of the remaining rainforest extent and support considerable biodiversity, and so are highly important for conservation. However, our data also indicate that such forests are potentially more fragile than their old growth counterparts, and so efforts should be made to reduce future environmental perturbations where possible.

## Acknowledgements

This study was funded by the UK Natural Environment Research Council to SJR, OL and MJS (under the Human-Modified Tropical Forests programme, NE/K016407/1; http://lombok.nerc-hmtf.info/), a Royal Society grant (RG130793) to ELC, and a Bat Conservation International grant to DRHB. We used Queen Mary’s Apocrita HPC facility, supported by QMUL Research-IT (http://doi.org/10.5281/zenodo.438045).

For assistance with data collection we thank Jamiluddin Jami, Arnold James, Mohd. Mustamin, Ampat Siliwong, Sabidee Mohd. Rizan, Najmuddin Jamal, Genevieve Durocher and Anne Seltmann. We thank Henry Bernard, Eleanor Slade and members of the LOMBOK consortium for facilitating research in Sabah, and we are grateful to the Sabah Biodiversity Council (licence numbers in Supplementary Information 10)

We thank Steven Le Comber, Hernani Oliveira, Joshua Potter, Sandra Álvarez Carretero and Kim Warren for their analytical assistance, and Mark Brown, Darren Evans and Talya Hackett for their comments on an earlier version of this manuscript. We thank Richard Sebastian and Omar Khalilur Rahman for their assistance with translating the manuscript abstract, and Craig Palmer for graphical assistance.

