## Supplementary Information for "Altered structure and stability of bat-prey interaction networks in logged tropical forests revealed by metabarcoding"

### Supplementary information 1: field work

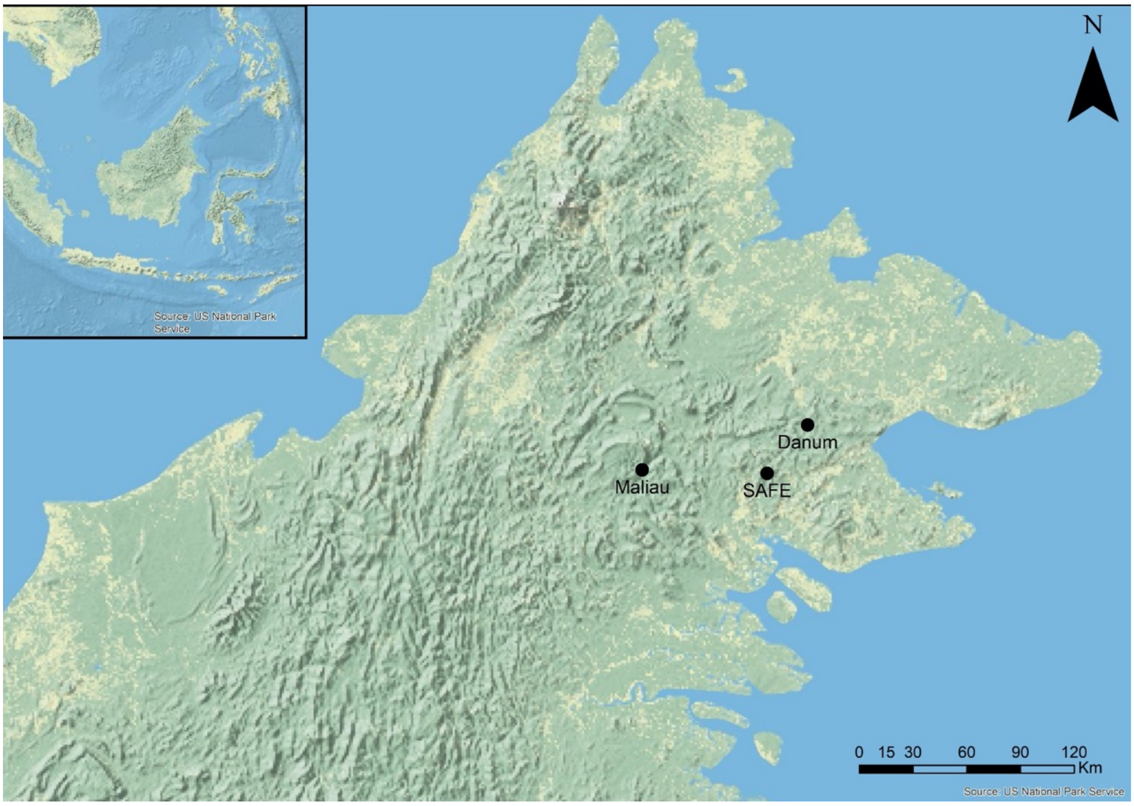


**Locations of study sites in Sabah, Malaysia.**

We sampled bats using six harp traps per night at three lowland tropical rainforest sites within Sabah, Malaysia, each <500m above sea level and with largely unseasonal climate. Two sites (Danum Valley and Maliau Basin) are primary rainforest, and the final site (the Stability of Altered Forest Ecosystems Project) have been subject to substantial anthropogenic disturbance.

With the exception of pregnant, lactating or juvenile individuals (which were released immediately after capture), bats were retained for up to 12 hours, usually around 10 hours, to allow sufficient time for defecation following the morning/evening's feeding activity. Sample sites were often substantial distances from the research camp, and so bats were placed in individual bags and transported to camp so they could be kept in quiet, humid conditions to minimise stress.

- The Danum Valley Conservation Area (hereafter 'Danum') is a 438 km$^2$ region of undisturbed primary rainforest in Sabah (Reynolds, Payne, Sinun, Mosigil, & Walsh, 2011), where there has been little logging or hunting in historic times. It is designated as a fully protected forest reserve. All of my sampling was in the forest <1.2 km from Danum Valley Field Centre. Traps were erected in 2016 for ten nights in a 21 night period and 2017 for ten nights in a 12 night period.

- The Maliau Basin Conservation Area (hereafter 'Maliau') is a 588 $km^2$ fully protected forest reserve made up of lowland and hill forest, most of which has neither been logged nor inhabited in historical times. WE sampled at Maliau Basin on the Seraya trail, an area of primary lowland rainforest. Traps were erected in 2016 and 2017 for ten nights in a 16 night period.

- The Stability of Altered Forest Ecosystems Project (hereafter 'SAFE') is a large ecological experiment where an area of degraded forest is being converted to oil palm plantation, with fragments of forest and riverine buffers being retained for scientific study (Ewers et al., 2011). The site has been subjected to disturbance since approximately March 2015, with areas of forest outside of designated fragments being salvage-logged. At the time that WE completed my sampling in July 2017, the forest was still in the process of conversion; thus, the study area was composed of experimental forest areas located within a matrix of highly degraded forest and shrubby clearings. WE sampled in the blocks 'LFE', 'B' and 'C', within the Ulu Segama Forest Reserve and Kalabakan area, during 2015, 2016 and 2017. Each of the blocks was sampled for a 5-night period, and then resampled at least 5 weeks later.

Total number of captures and samples collected

| | *Number captured* | *Number of samples obtained* | *Rate* |
|---|---|---|---|
| *Balionycteris maculata* | 5 | 1 | 0.2 |
| *Emballonura monticola* | 2 | 1 | 0.5 |
| *Hesperoptenus blanfordi* | 19 | 10 | 0.526 |
| *Hipposideros ater* | 5 | 1 | 0.2 |
| *Hipposideros bicolor* | 10 | 6 | 0.6 |
| *Hipposideros cervinus* | 1147 | 493 | 0.43 |
| *Hipposideros diadema* | 44 | 24 | 0.545 |
| *Hipposideros dyacorum* | 57 | 25 | 0.439 |
| *Hipposideros galeritus* | 1 | 1 | 1 |
| *Hipposideros ridleyi* | 39 | 24 | 0.615 |
| *Kerivoula hardwickii* | 99 | 33 | 0.333 |
| *Kerivoula intermedia* | 273 | 97 | 0.355 |
| *Kerivoula lenis* | 3 | 3 | 1 |
| *Kerivoula minuta* | 12 | 4 | 0.333 |
| *Kerivoula papillosa* | 103 | 40 | 0.388 |
| *Kerivoula pellucida* | 29 | 11 | 0.379 |
| *Megaderma spasma* | 5 | 3 | 0.6 |
| *Murina aenea* | 1 | 1 | 1 |
| *Murina cyclotis* | 4 | 2 | 0.5 |
| *Murina peninsularis* | 4 | 4 | 1 |
| *Murina rozendaali* | 5 | 3 | 0.6 |
| *Murina suilla* | 16 | 9 | 0.562 |
| *Myotis muricola* | 3 | 3 | 1 |
| *Myotis ridleyi* | 1 | 1 | 1 |
| *Nycteris tragata* | 7 | 2 | 0.286 |
| *Phoniscus atrox* | 3 | 1 | 0.333 |

The number of captures and samples obtained for the bat species used to make the ecological networks, and the defecation rate per species.

| | Number captured | Number of samples obtained | Rate |
|---|---|---|---|
| *Hipposideros cervinus* | 1147 | 493 | 0.43 |
| *Hipposideros diadema* | 44 | 24 | 0.545 |
| *Hipposideros dyacorum* | 57 | 25 | 0.439 |
| *Kerivoula hardwickii* | 99 | 33 | 0.333 |
| *Kerivoula intermedia* | 273 | 97 | 0.355 |
| *Rhinolophus borneensis* | 103 | 40 | 0.388 |
| *Rhinolophus sedulus* | 58 | 37 | 0.638 |
| *Rhinolophus trifoliatus* | 81 | 34 | 0.42 |

### Supplementary information 2: analysis

#### Laboratory work

WE undertook DNA extraction and PCR using the protocol outlined in Chapter 2. Quality-control on amplicons was then performed using a DNA D1000 TapeStation (Agilent Technologies), and quantified using a QuBit dsDNA HS Assay Kit (Invitrogen, Life Technologies). Finally, sequencing took place on three 96-well plates, which were run on an Illumina MiSeq at the Genome Centre, London (UK). Sequencing was performed bi-directionally with Fluidigm indexes following manufacturer's instructions.

#### Bioinformatics pipeline

WE assembled forward and reverse sequence reads into contigs using mothur (Schloss et al., 2009), and then removed forward and reverse primers using the galaxy web platform on the public server at usegalaxy.org (Afgan et al., 2016), as per the protocol in Chapter 2. A custom python script was used to remove any haplotypes either represented by a single sequence or falling outside of a length of 155-159bp. WE then generated subsequent datasets using MOTU clustering thresholds at ranges 91-98% similarity, using the Uclust algorithm (Edgar, 2010) as implemented in the QIIME platform (Caporaso, Kuczynski, Stombaugh, Bittinger, & Bushman, 2010). Representative sequences for each MOTU per clustering level were then compared to one another using BLAST+ (Camacho et al., 2009), with the resulting data being reduced in LULU (Frøslev et al., 2017) for quality control. All resulting bat-MOTU adjacency lists were then transformed into adjacency matrices using a custom perl script. These matrices were then split into multiple binary adjacency matrices by site, where *aij* denotes the consumption of MOTU *j* by bat individual *i*. Networks were created by pooling samples from multiple years. All bioinformatic and statistical steps are recorded at https://github.com/hemprichbennett/bat-diet.

**Supplementary information 3. The number of MOTUs consumed by bats at each site, as observed at each MOTU clustering level.**

| Clustering level | **Danum** | **Maliau** | **SAFE** |
|---|---|---|---|
| 91 | 1329 | 920 | 1205 |
| 92 | 1373 | 884 | 1243 |
| 93 | 1369 | 875 | 1237 |
| 94 | 1378 | 876 | 1203 |
| 95 | 1388 | 842 | 1207 |
| 96 | 1401 | 837 | 1224 |
| 97 | 1449 | 841 | 1256 |
| 98 | 1544 | 873 | 1301 |

#### Supplementary information 4. Closeness centrality of each bat species studied

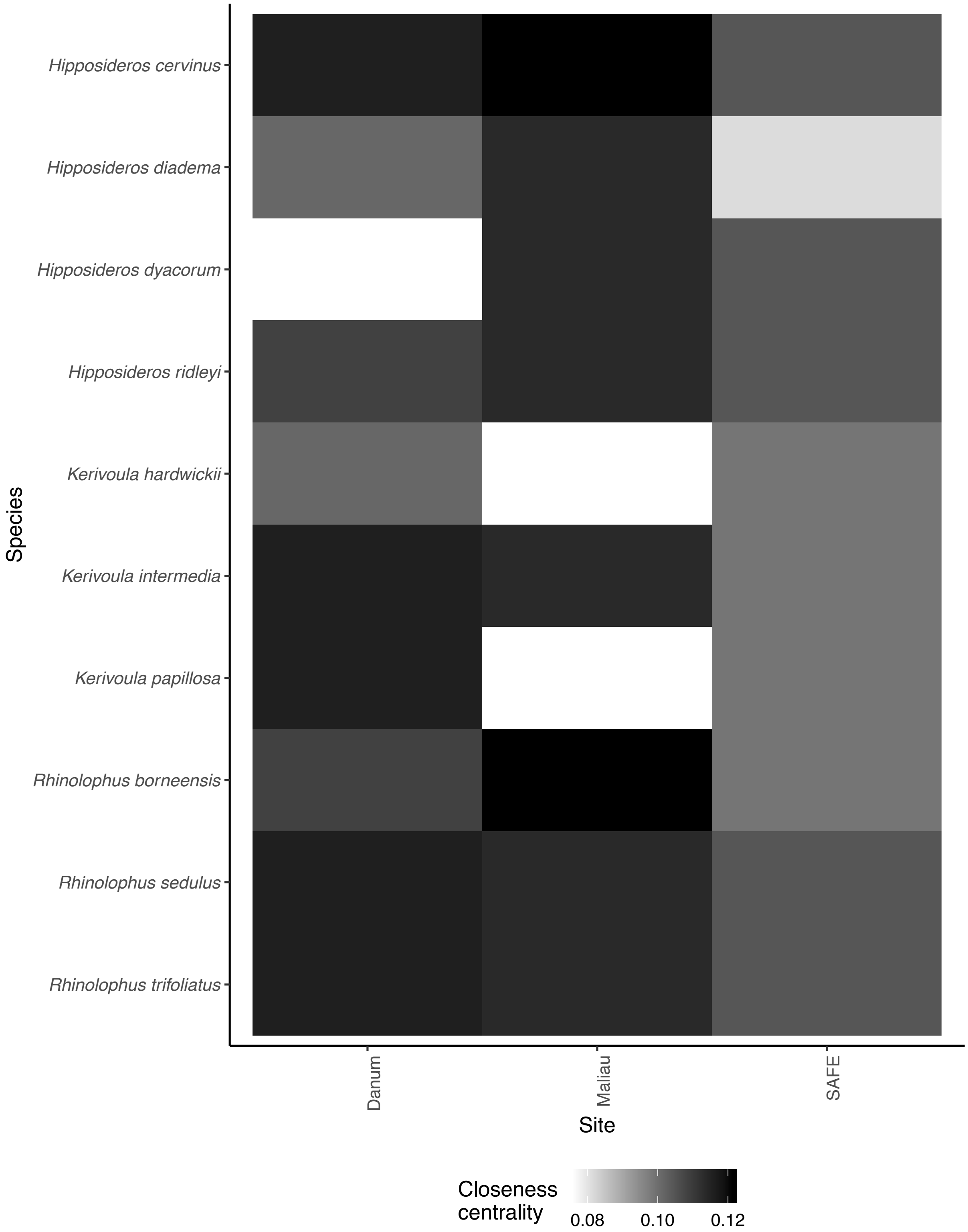

#### Supplementary information 5. Betweenness centrality of each bat species studied

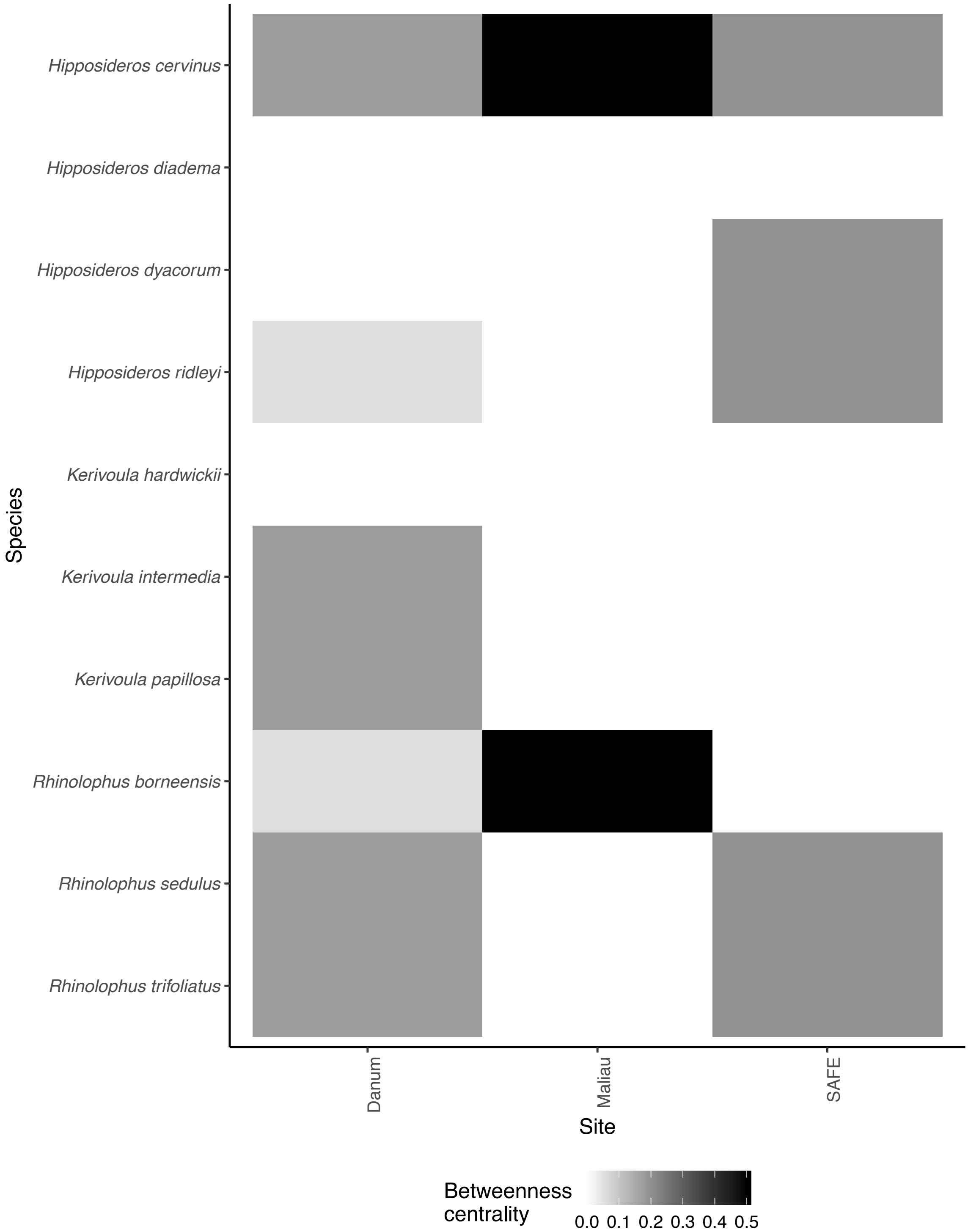

#### Supplementary information 6. Rankings of the influence of each bat species on a given network-level metric.

1 is the ranking of highest influence, 10 is the lowest. White tiles denote a bat species not included in the network for that site, due to lack of captures.

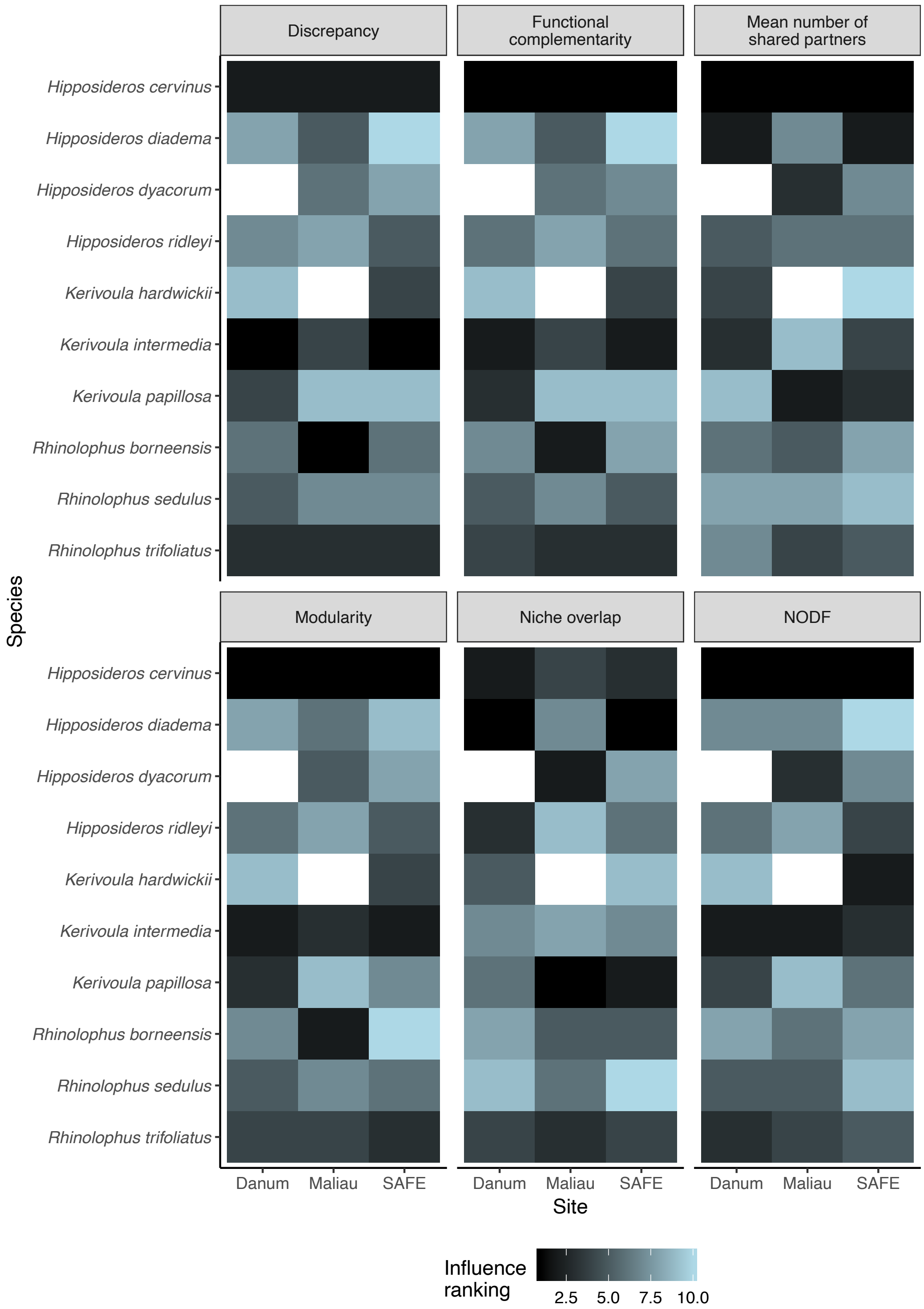

**Supplementary information 7. Alterations in observed metric values when rarefying networks**

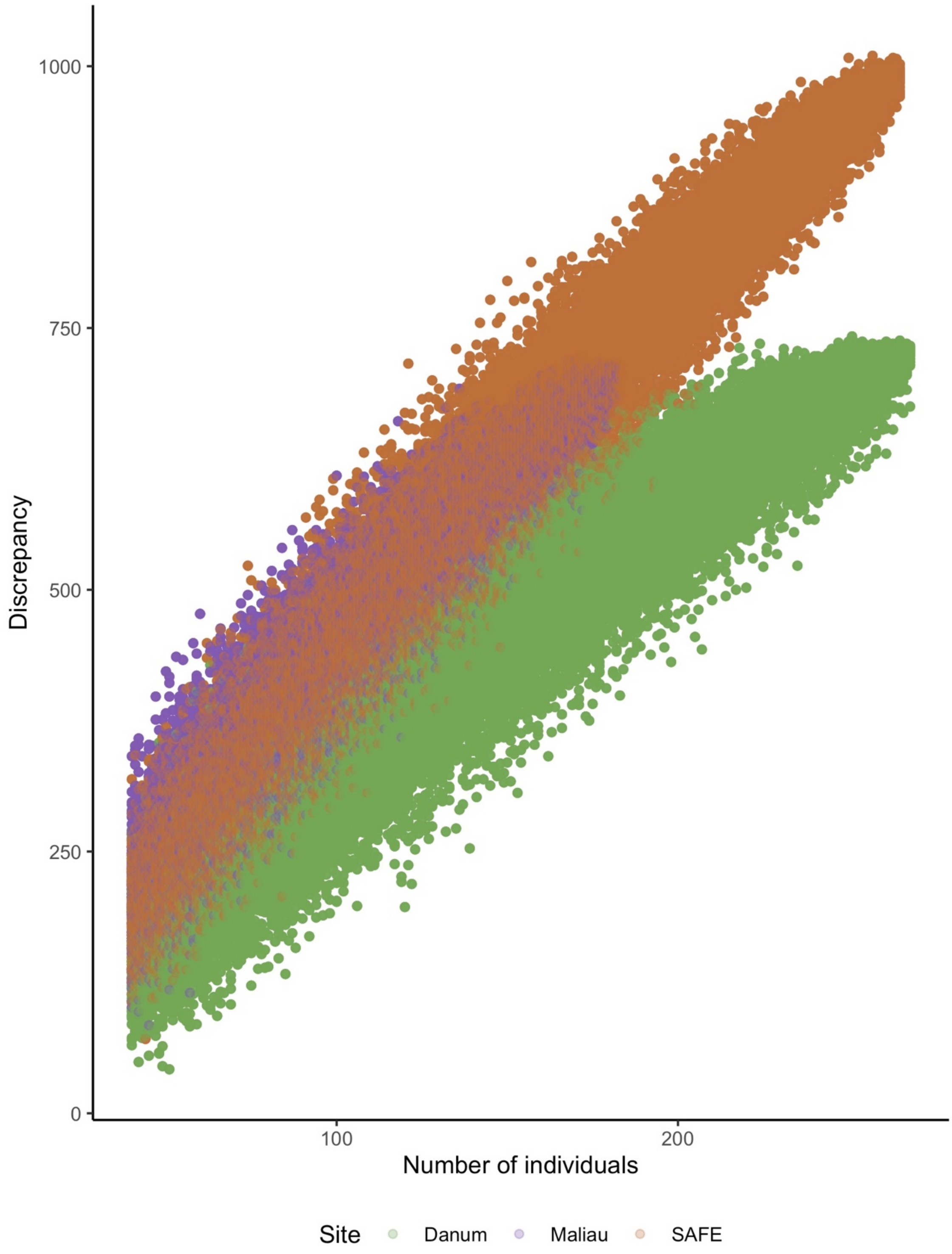

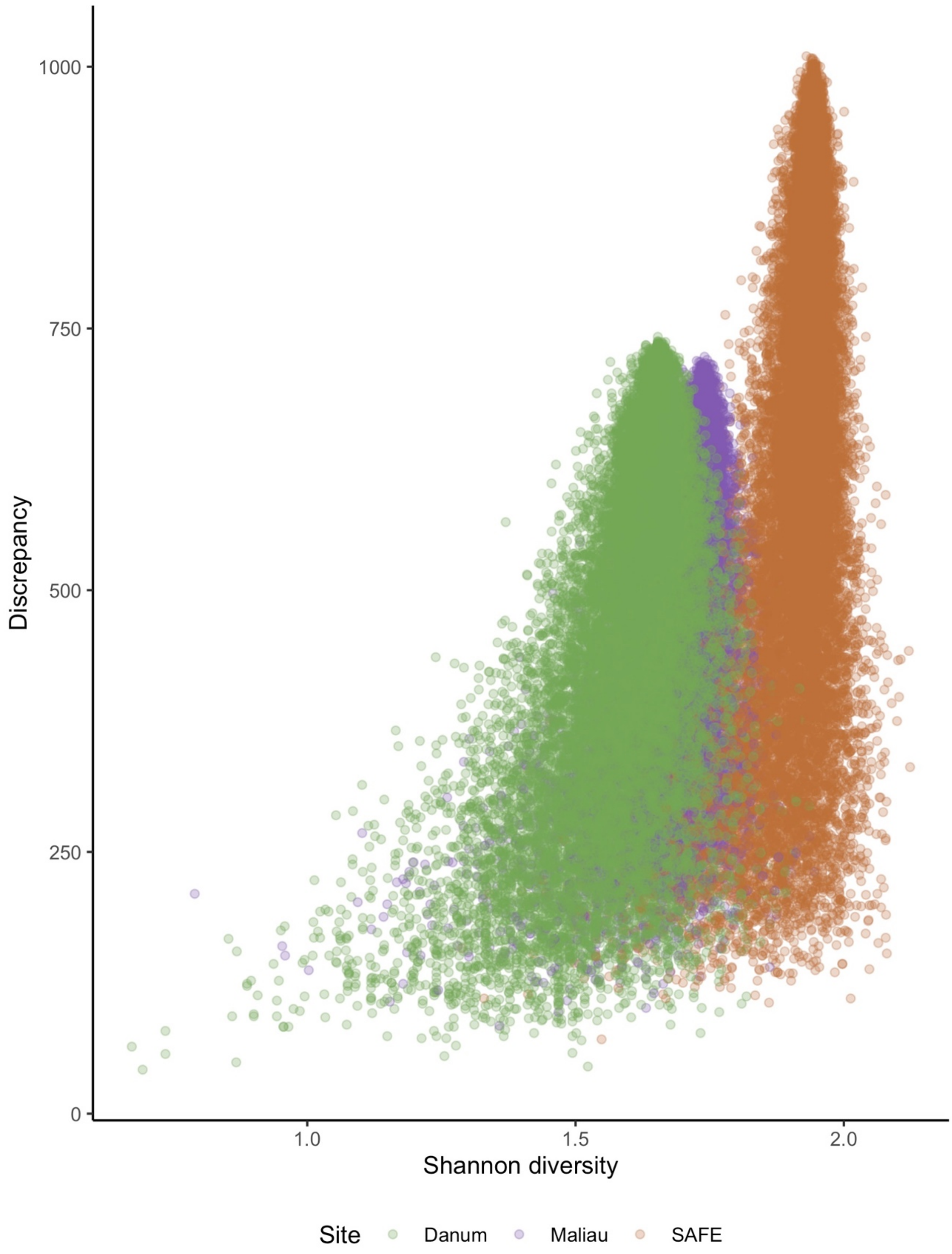

Discrepancy
1000
750
500
250
0
1.0
1.5
2.0
Shannon diversity
Site
Danum
Maliau
SAFE

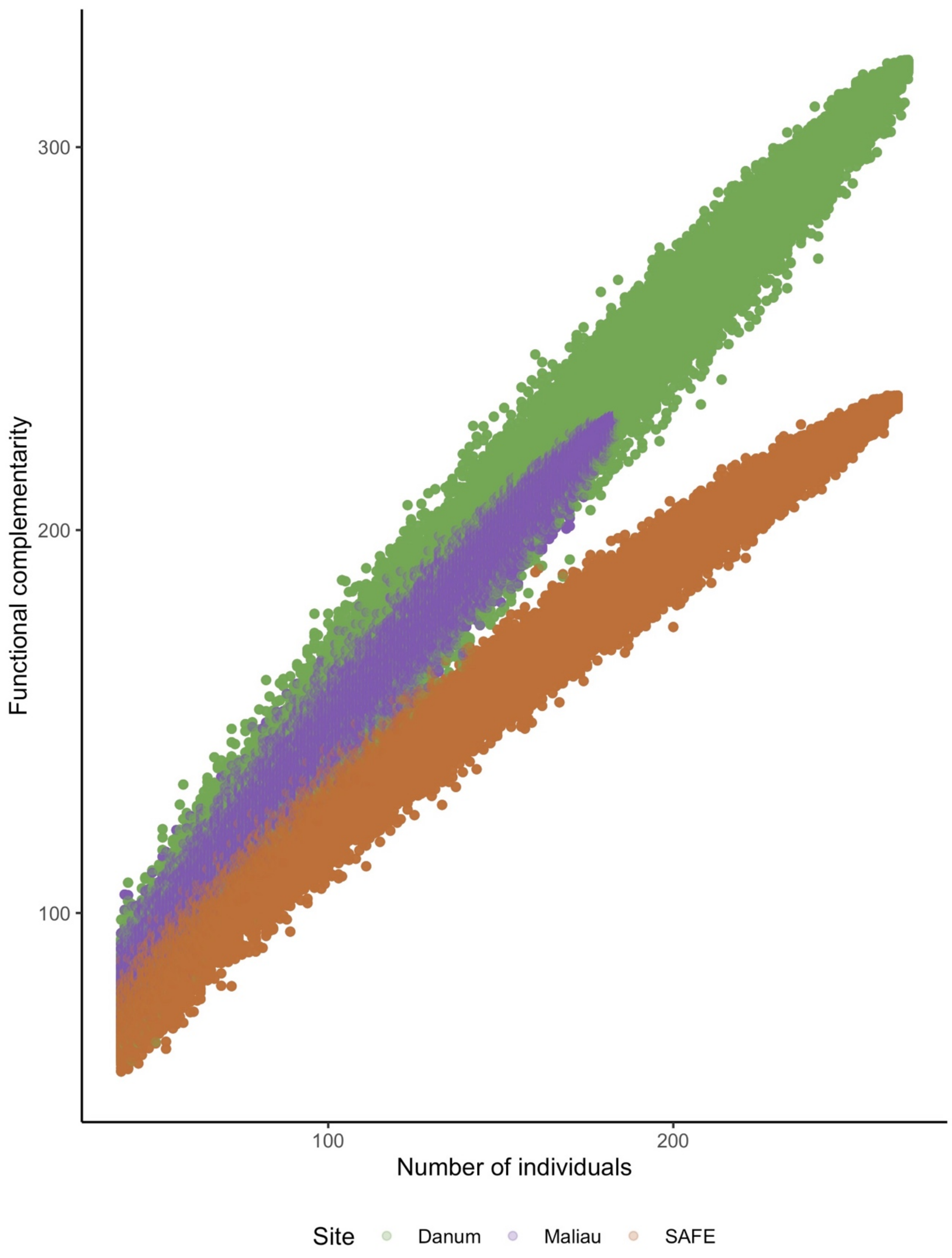

Functional complementarity
300
200
100
100
200
Number of individuals
Site
Danum
Maliau
SAFE

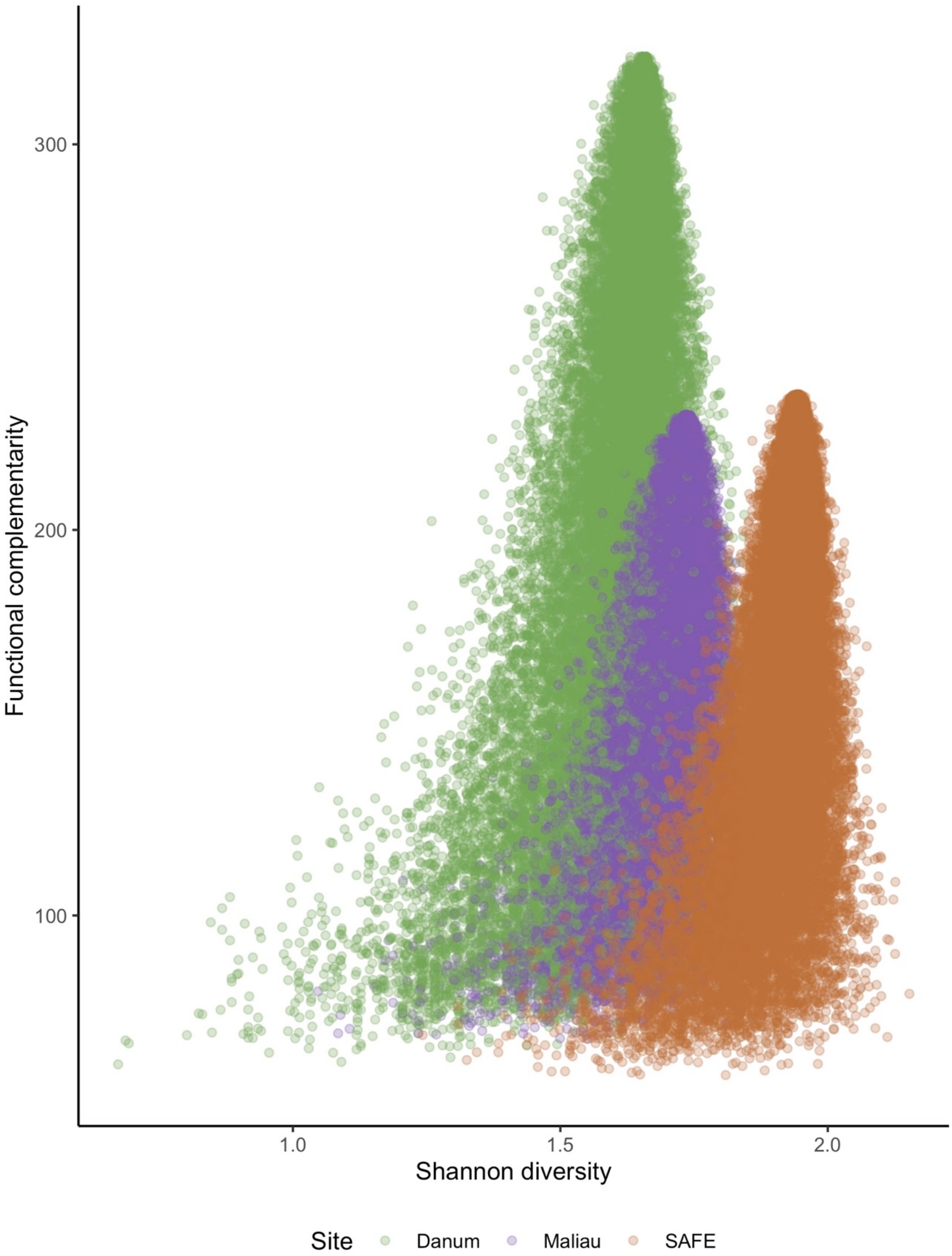

Functional complementarity
300
200
100
1.0
1.5
2.0
Shannon diversity
Site
Danum
Maliau
SAFE

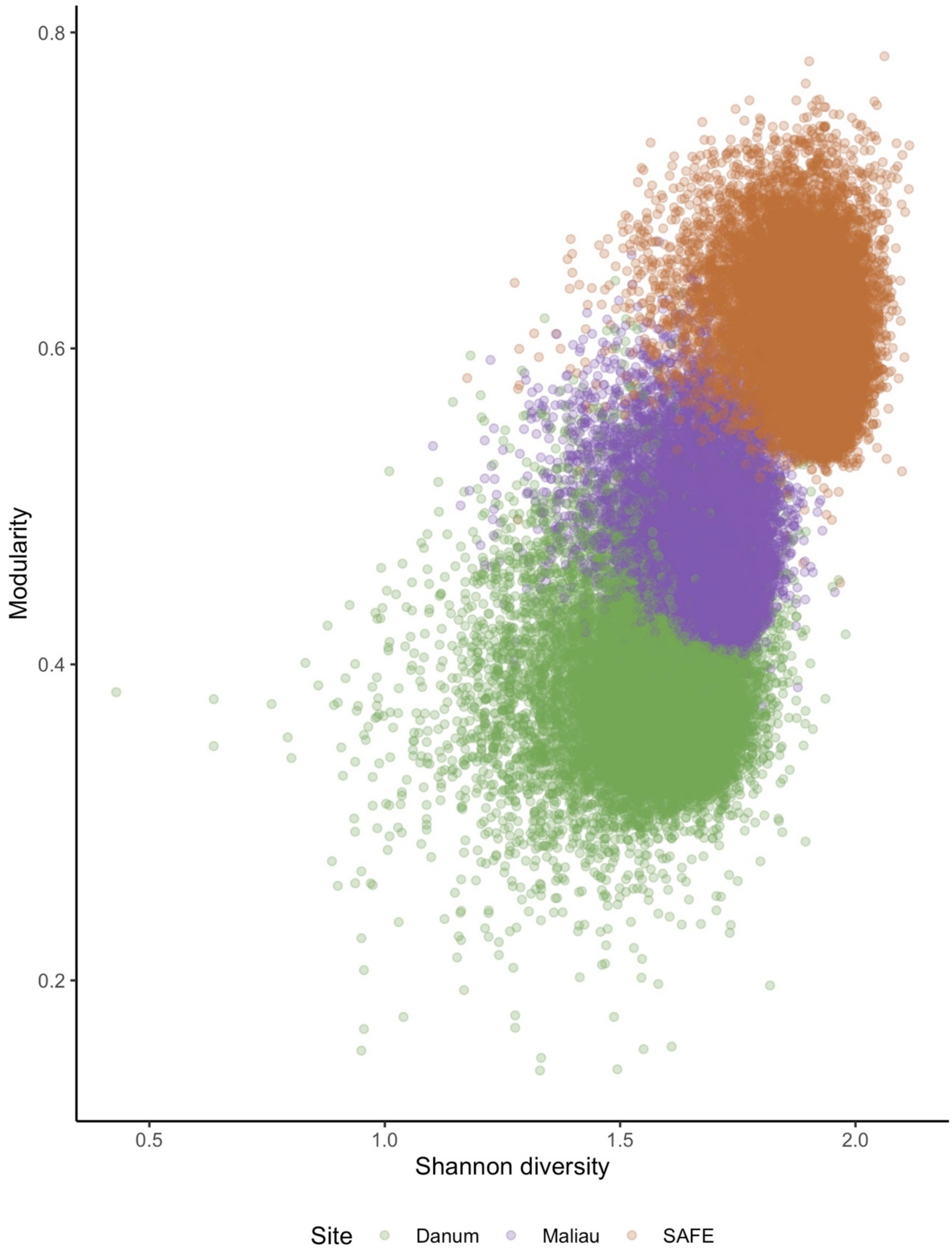

0.8
0.6
0.4
0.2
Modularity
0.5
1.0
1.5
2.0
Shannon diversity
Site
Danum
Maliau
SAFE

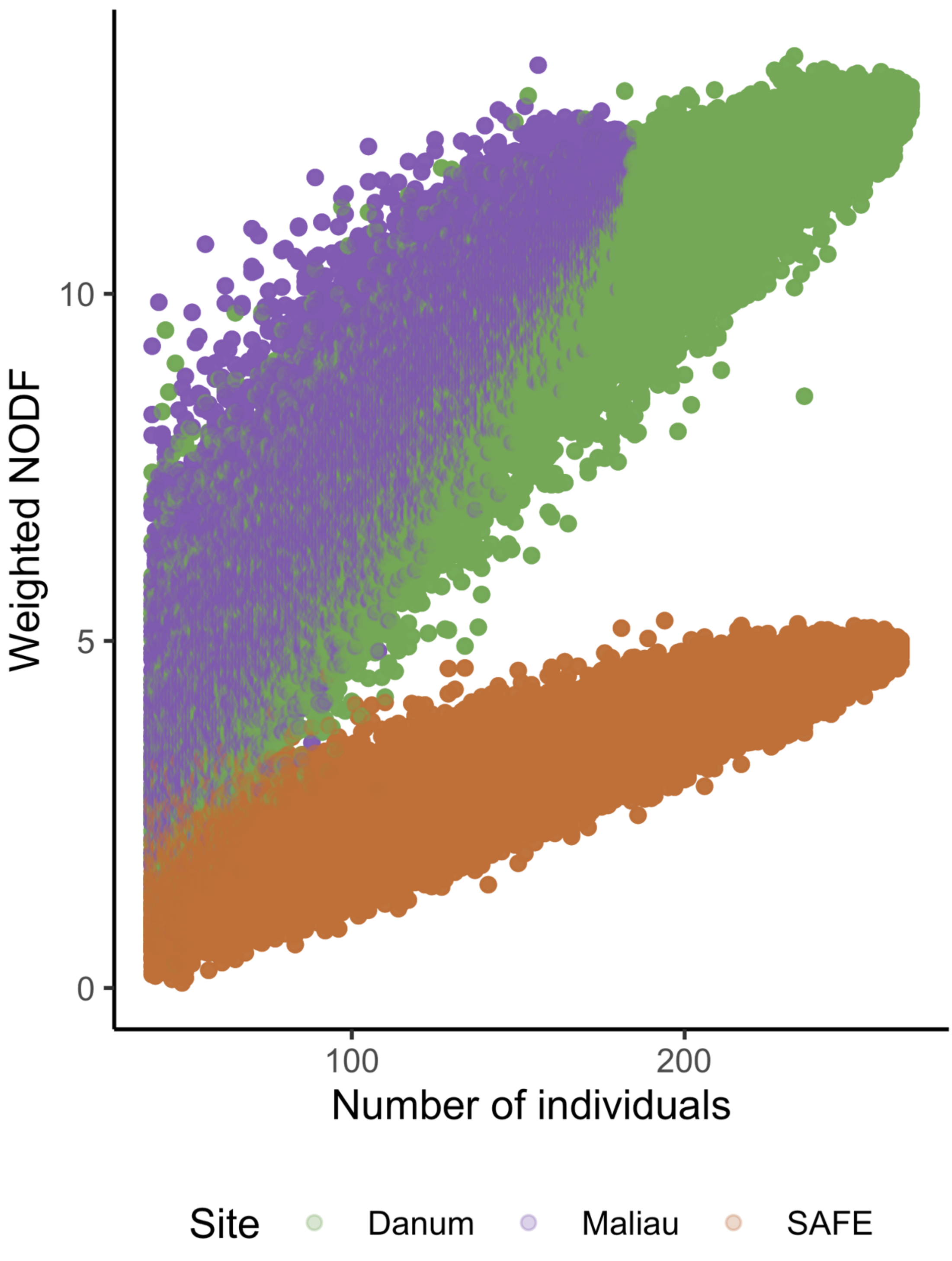
Weighted NODF
0
5
10
100
200
Number of individuals
Site
Danum
Maliau
SAFE

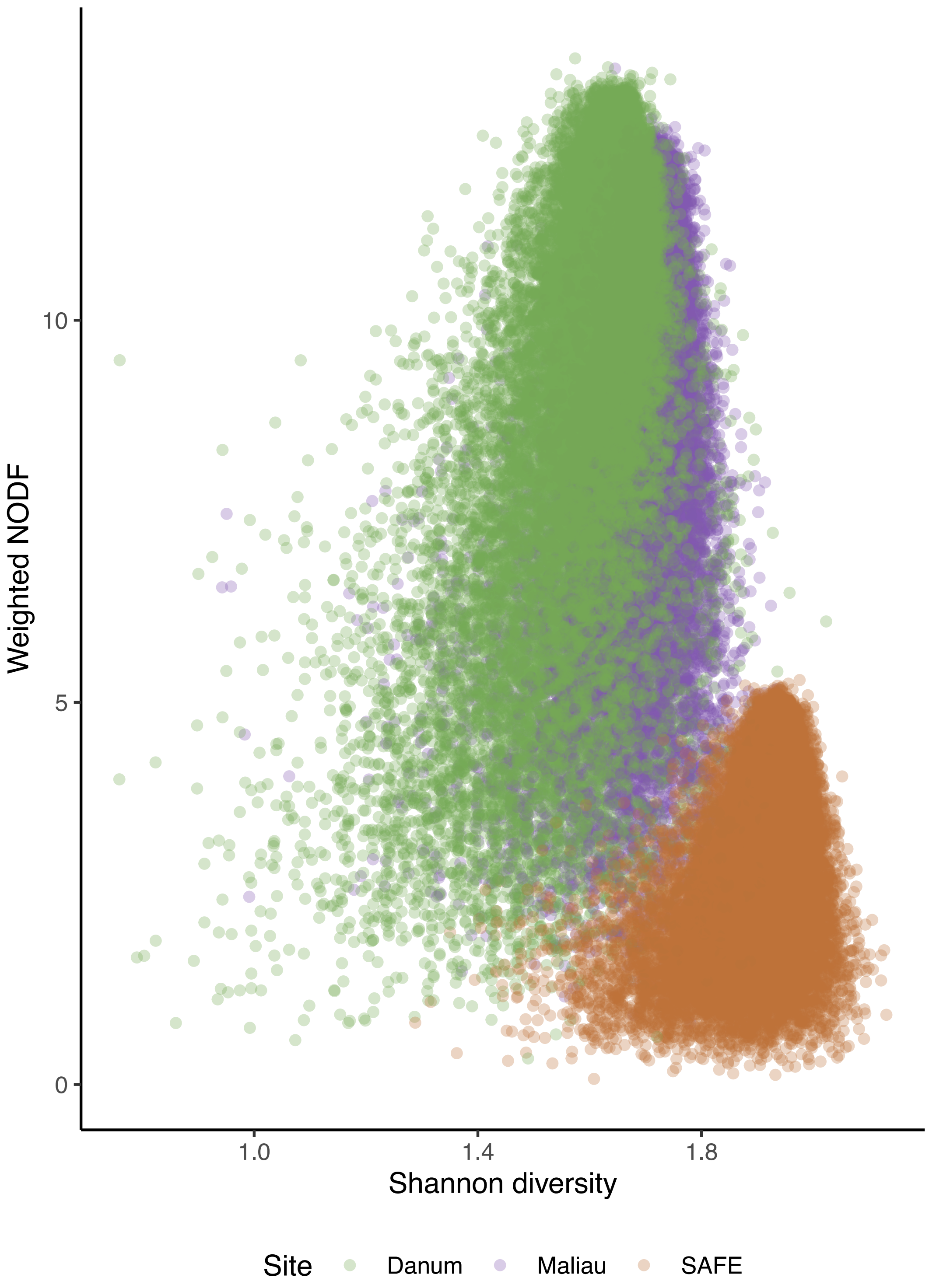

Weighted NODF
10
5
0
1.0
1.4
1.8
Shannon diversity
Site
Danum
Maliau
SAFE

### Supplementary information 8. Family-level diets of all bats sequenced

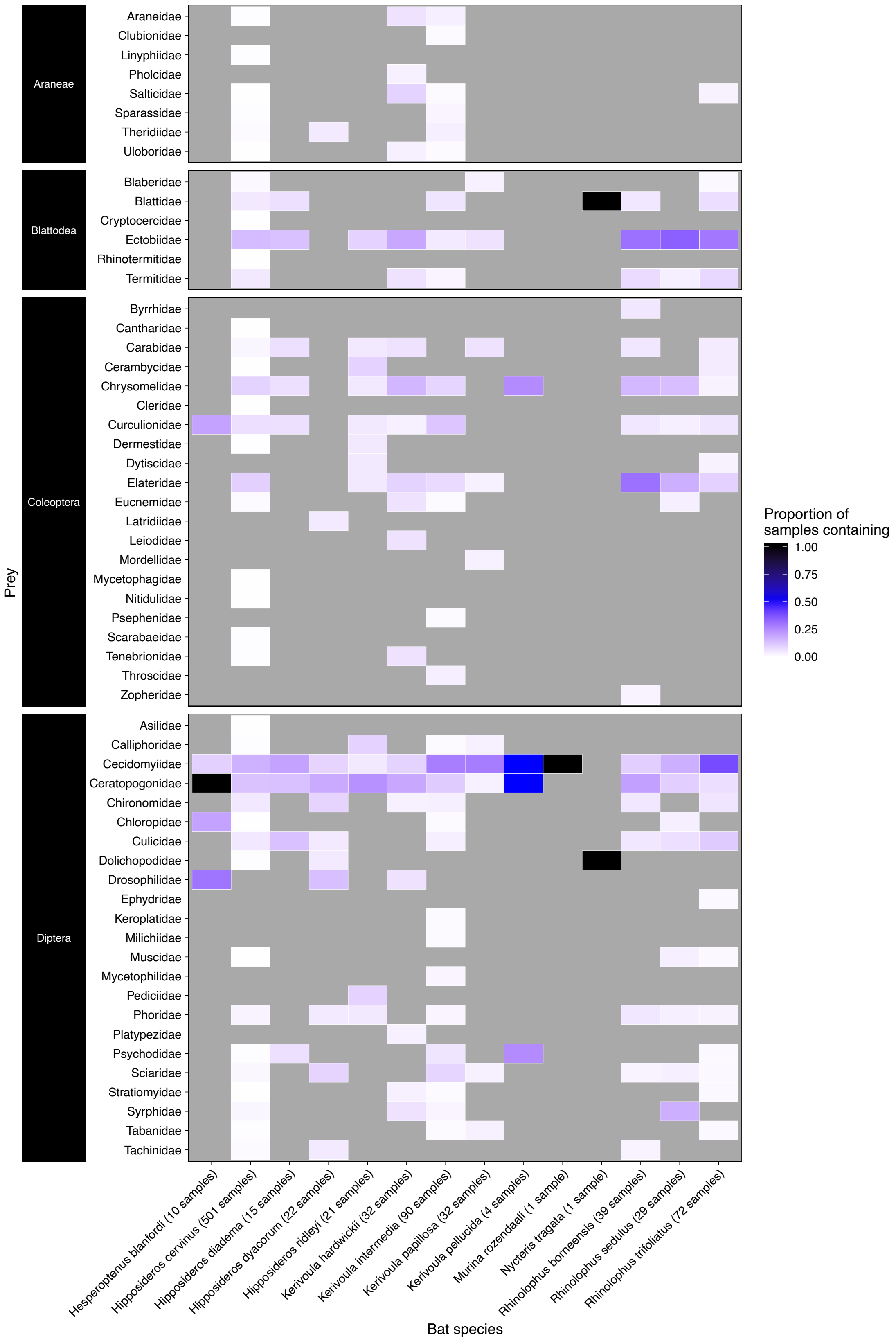

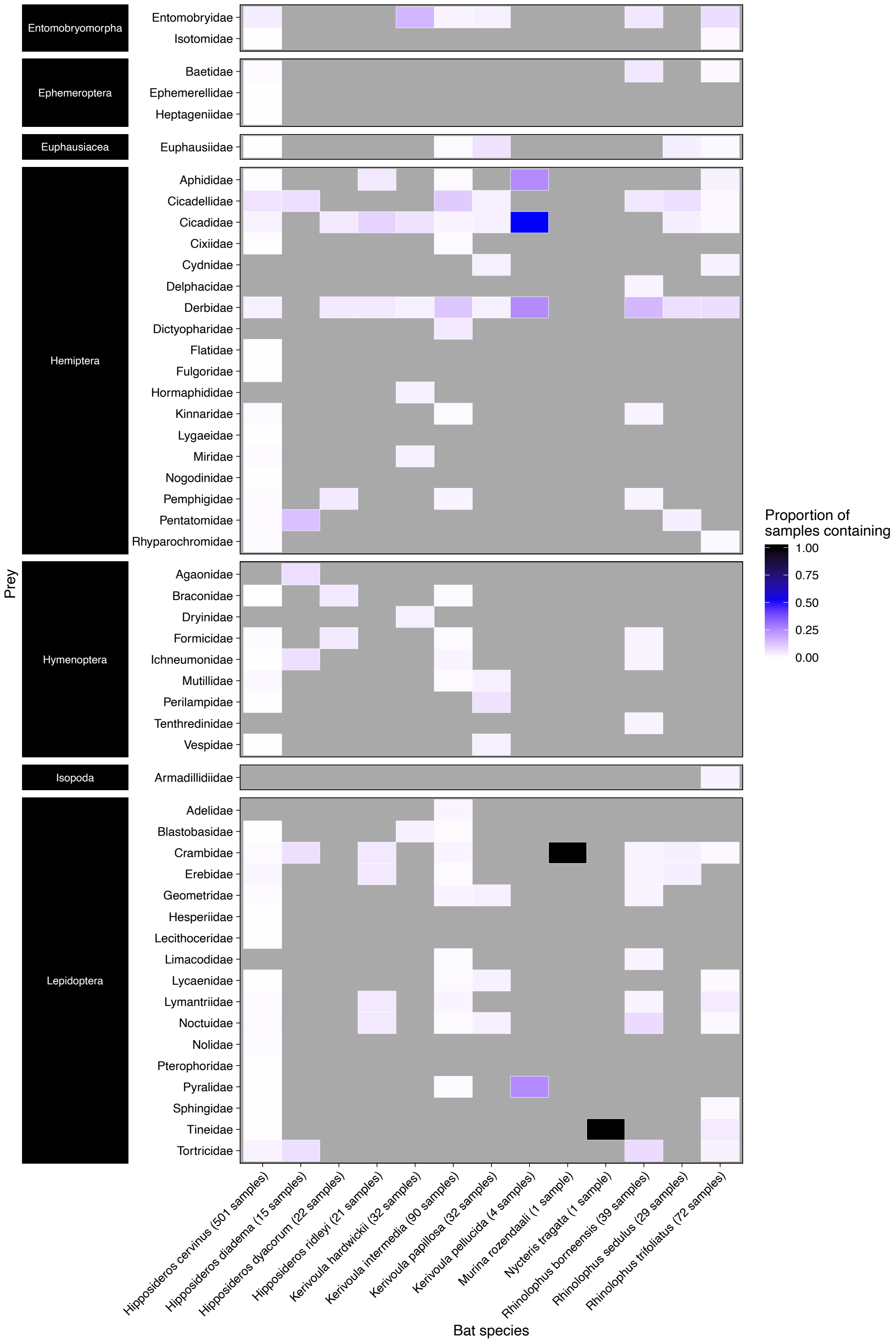
Entomobryomorpha
Entomobryidae
Isotomidae
Ephemeroptera
Baetidae
Ephemerellidae
Heptageniidae
Euphausiacea
Euphausiidae
Hemiptera
Aphididae
Cicadellidae
Cicadidae
Cixiidae
Cydnidae
Delphacidae
Derbidae
Dictyopharidae
Flatidae
Fulgoridae
Hormaphididae
Kinnaridae
Lygaeidae
Miridae
Nogodinidae
Pemphigidae
Pentatomidae
Rhyparochromidae
Hymenoptera
Agaonidae
Braconidae
Dryinidae
Formicidae
Ichneumonidae
Mutillidae
Perilampidae
Tenthredinidae
Vespidae
Isopoda
Armadillidiidae
Lepidoptera
Adelidae
Blastobasidae
Crambidae
Erebidae
Geometridae
Hesperiidae
Lecithoceridae
Limacodidae
Lycaenidae
Lymantriidae
Noctuidae
Nolidae
Pterophoridae
Pyralidae
Sphingidae
Tineidae
Tortricidae
Prey
Hipposideros cervinus (501 samples)
Hipposideros diadema (15 samples)
Hipposideros dyacorum (22 samples)
Hipposideros ridleyi (21 samples)
Kerivoula hardwickii (32 samples)
Kerivoula intermedia (90 samples)
Kerivoula papillosa (32 samples)
Kerivoula pellucida (4 samples)
Murina rozendaali (1 sample)
Nycteris tragata (1 sample)
Rhinolophus borneensis (39 samples)
Rhinolophus sedulus (29 samples)
Rhinolophus trifoliatus (72 samples)
Bat species
Proportion of
samples containing
1.00
0.75
0.50
0.25
0.00

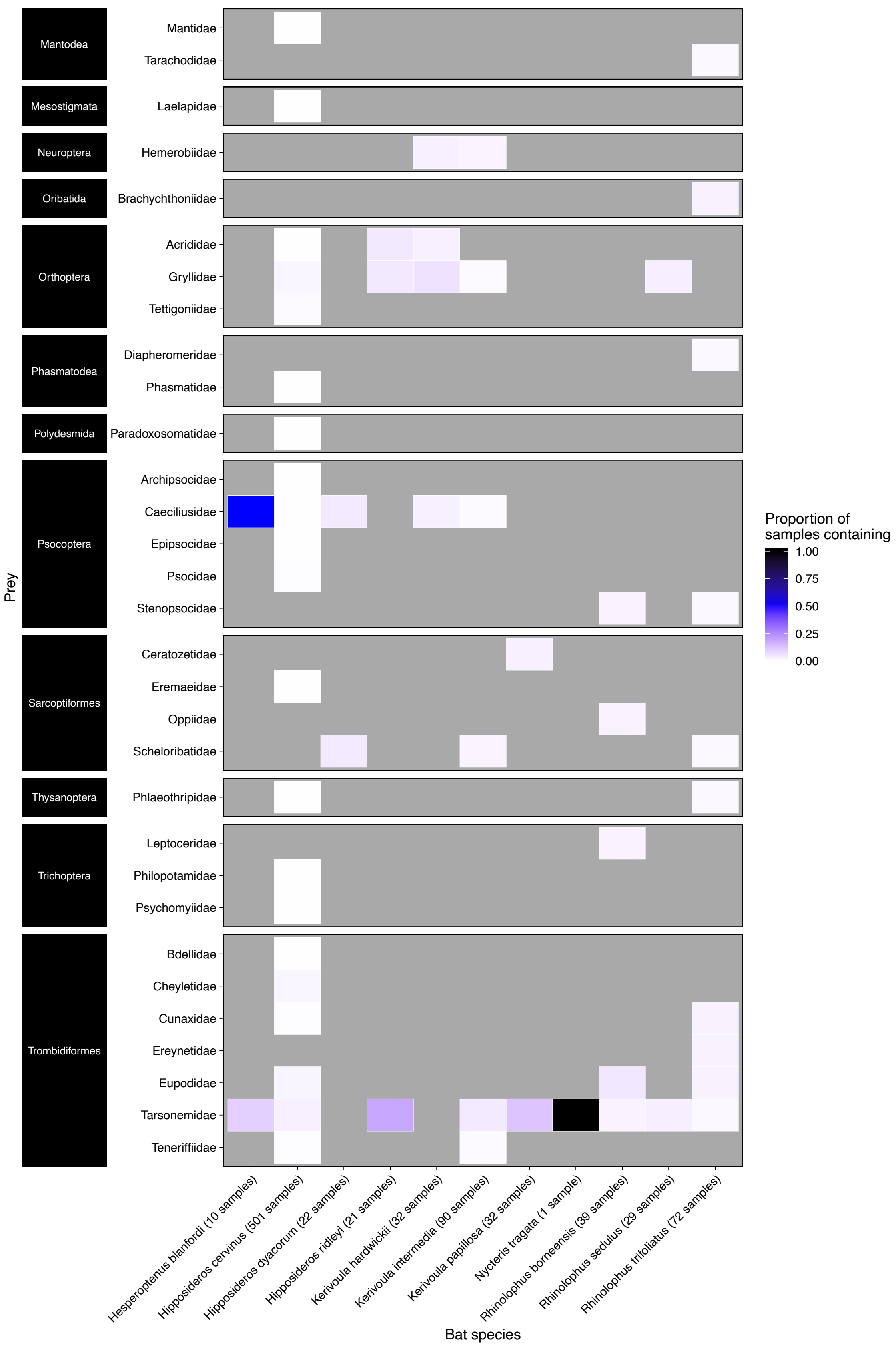

Mantodea
Mantidae
Tarachodidae
Mesostigmata
Laelapidae
Neuroptera
Hemerobiidae
Oribatida
Brachychthoniidae
Orthoptera
Acrididae
Gryllidae
Tettigoniidae
Phasmatodea
Diapheromeridae
Phasmatidae
Polydesmida
Paradoxosomatidae
Psocoptera
Archipsocidae
Caeciliusidae
Epipsocidae
Psocidae
Stenopsocidae
Sarcoptiformes
Ceratozetidae
Eremaeidae
Oppiidae
Scheloribatidae
Thysanoptera
Phlaeothripidae
Trichoptera
Leptoceridae
Philopotamidae
Psychomyiidae
Trombidiformes
Bdellidae
Cheyletidae
Cunaxidae
Ereynetidae
Eupodidae
Tarsonemidae
Teneriffiidae
Prey
Hesperoptenus blanfordi (10 samples)
Hipposideros cervinus (501 samples)
Hipposideros dyacorum (22 samples)
Hipposideros ridleyi (21 samples)
Kerivoula hardwickii (32 samples)
Kerivoula intermedia (90 samples)
Kerivoula papillosa (32 samples)
Nycteris tragata (1 sample)
Rhinolophus borneensis (39 samples)
Rhinolophus sedulus (29 samples)
Rhinolophus trifoliatus (72 samples)
Bat species
Proportion of
samples containing
1.00
0.75
0.50
0.25
0.00

### Supplementary information 9 Dietary correlations

**Methods**

To identify if there are patterns of taxonomic Orders co-occurring within the diet of individual bats (e.g. bat individuals that feed on Coleoptera may be more likely to feed on Blattodea), we analysed the co-occurrence between taxonomic Orders within the diets of bats using Pearson's correlation coefficient, with each individual bat acting as a sample. This took place to identify potential significant correlations of prey consumption, and potential primer biases as the ZBJ primers used here have previously been shown to preferentially amplify some taxa (Alberdi, Aizpurua, Gilbert, & Bohmann, 2018). The analysis was restricted to bat individuals that had consumed more than five taxonomically-matched MOTUs, and to Orders of prey that had been consumed by over 20 bats; this approach reduced the biases arising from zero-values. The resulting data was visualised using the R package 'corrplot' (Wei & Simko, 2017).

**Results:**

Significant positive and negative correlations occurred between many taxonomic Orders (see Supplementary Information), with Hymenoptera and Orthoptera being the only Orders negatively correlated with multiple Orders, though correlations were weak. Despite reported primer bias towards Lepidoptera and Diptera (Alberdi et al., 2018), these Orders were each only negatively correlated with a single other Order, and these correlations were also weak.

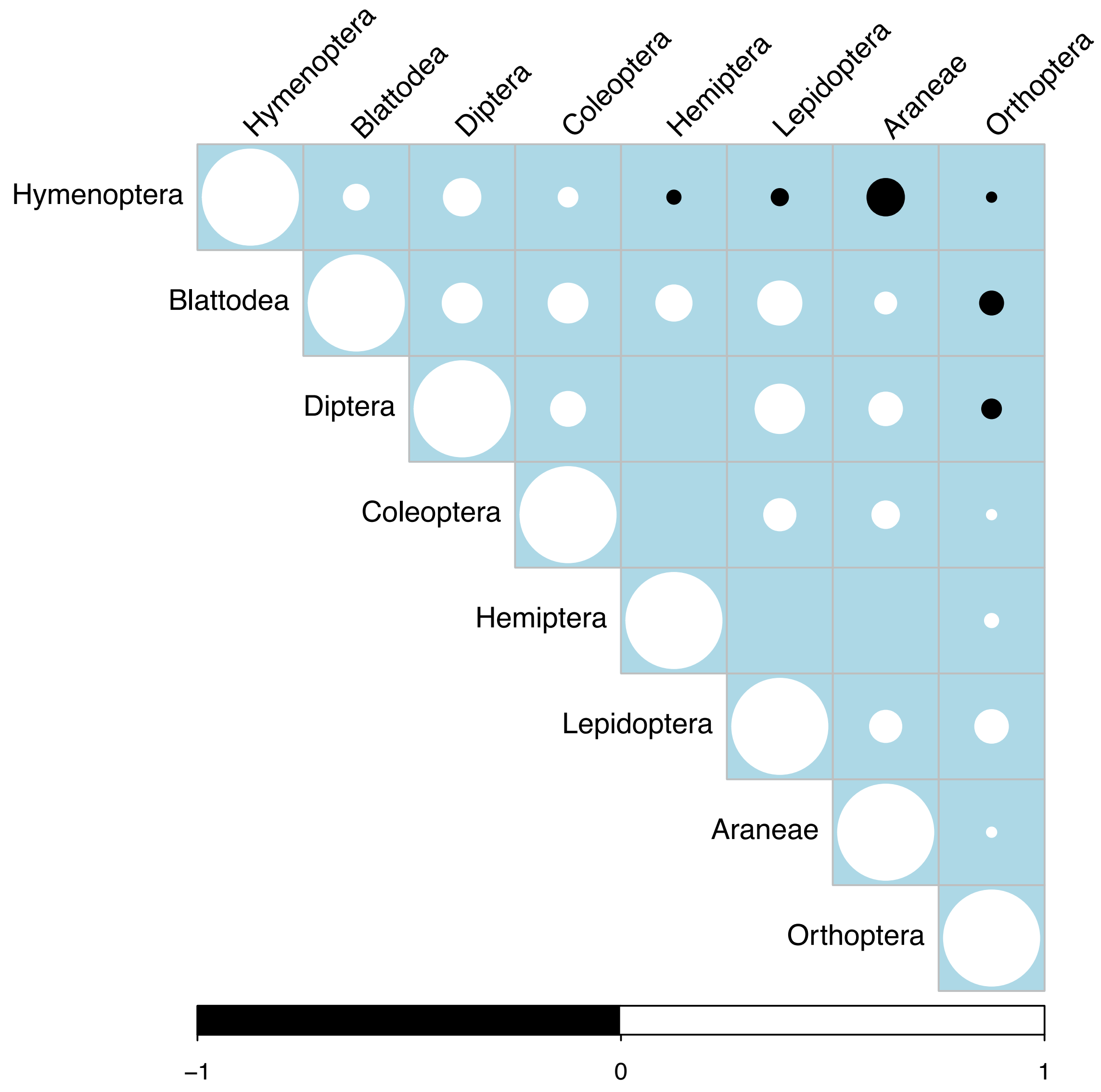


**Positive and negative correlations of orders consumed by all bats sequenced.** Only bats consuming more than 5 taxonomically identified MOTUs, and orders consumed by more than 20 bats were included. Circle size is proportional to the strength of the correlation, black circles are negative correlations, white circles are positive correlations. Empty locations within the grid show non-significant ($p>0.05$) correlations.

#### Supplementary information 10: licenses and permits for fieldwork

(access licenses JKM/MBS.1000- 2/2 (374), JKM/MBS.1000-2/2 JLD.4 (23), JKM/MBS.1000-2/2 JLD.4 (45), JKM/MBS.1000-2/2 JLD.4 (41), Access licenses JKM/MBS.1000-2/2 JLD.4 (46), JKM/MBS.1000-2/2 JLD.5 (123) and JKM/MBS.1000-2/2 JLD.5 (153), export licenses JKM/MBS.1000-2/3 JLD.2(55), JKM/MBS.1000-2/3 JLD.2 (95) and JKM/MBS.1000-2/3 JLD.3 (31). Danum Valley access permits (refs YS/DVMC/2015/221, YS/DVMC/2016/11, YS/DVMC/2015/222, YS/DVMC/2016/13, YS/DVMC/2017/42, YS/DVMC/2017/41) Maliau Basin access permits (refs YS/MBMC/2015/186, YS/MBMC/2016/23, YS/MBMC/2015/187, YS/MBMC/2016/25, YS/MBMC/2017/67 and YS/MBMC/2017/66).
